# TOB is an effector of the hippocampus-mediated acute stress response

**DOI:** 10.1101/2022.05.16.492218

**Authors:** Mohieldin Youssef, Hiro Taiyo Hamada, Esther Suk King Lai, Yuji Kiyama, Mohamed Eltabbal, Hiroshi Kiyonari, Kohei Nakano, Bernd Kuhn, Tadashi Yamamoto

## Abstract

Stress affects behavior and involves critical dynamic changes at multiple levels ranging from molecular pathways to neural circuits and behavior. Abnormalities at any of these levels lead to decreased stress resilience and pathological behavior. However, temporal modulation of molecular pathways underlying stress response remains poorly understood. Transducer of ErbB2.1, known as TOB, (TOB1) is involved in different physiological functions, including cellular stress and immediate response to stimulation. In this study, we investigated the role of TOB in the brain’s stress machinery at molecular, neural circuit, and behavioral levels. Interestingly, TOB protein levels increased after mice were exposed to acute stress. At the neural circuit level, functional magnetic resonance imaging (fMRI) suggested that intra-hippocampal and hippocampal-prefrontal connectivity were dysregulated in *Tob* knockout (*Tob*-KO) mice. Electrophysiological recordings in hippocampal slices showed increased postsynaptic AMPAR-mediated neurotransmission, accompanied by decreased GABA neurotransmission and subsequently altered Excitatory/Inhibitory balance after *Tob* deletion. At the behavioral level, *Tob*-KO mice show abnormal, hippocampus-dependent, contextual fear conditioning and extinction, and depression-like behaviors. On the other hand, increased anxiety observed in *Tob*-KO mice is hippocampus-independent. At the molecular level, we observed decreased stress-induced LCN2 expression and ERK phosphorylation, as well as increased MKP-1 expression. This study suggests that TOB serves as an important modulator in hippocampal stress signaling machinery. In summary, we show a molecular pathway and neural circuit mechanism by which TOB deletion contributes to expression of pathological stress-related behavior.

## Introduction

On a daily basis, we encounter stressful events to which our bodies generate different responses and store memories to cope with future occurrences. The brain utilizes several mechanisms to cope with psychological stress, and defects in such mechanisms or exposure to excessive stress can increase individual vulnerability to neuropsychiatric disorders like depression and post-traumatic stress disorder (PTSD)^1^. Strikingly, it is estimated that 50% of adults have experienced a traumatic event during their lifetimes. Therefore, it is imperative that we investigate mechanisms that underlie stress responses and identify potential therapeutic targets coordinating stress resilience^2,3^.

The stress coping response is orchestrated at various intercalated layers which include brain connectivity, neuronal activity, molecular signaling, and resulting behavior^4^. Any change in stress resilience mechanisms can induce psychiatric consequences, such as increased fear, anxiety, and depression. Such behaviors are controlled by neuronal circuits governing emotional and fight-flight responses, like the hippocampus, prefrontal cortex, amygdala, and hypothalamus^5^. fMRI is currently the most advanced, non-invasive method to map dynamic changes in brain circuits that regulate stress coping^6, 7^. In response to stress, abnormal neuronal circuit remodeling may occur, leading to altered brain connectivity. Several molecules have been implicated in these remodeling events, like lipocalin-2 (LCN2) and corticotrophin-releasing factor (CRF) ^8^. The Hypothalamic-Pituitary Adrenal (HPA) axis is a hormonal signaling pathway that is moderately activated to elicit adaptation to induced stress at molecular, cellular, physiological, and behavioral levels^9^. At the molecular level, acute stress induces transcriptional and translation responses in order to cope with stress^10, 11^. This transient change in molecular signaling is believed to have neuronal protective functions^12^. Our knowledge of the hippocampal molecular stress machinery is limited; therefore, there are continuing efforts to identify genes that function in stress coping responses^13^. Interestingly, several molecules with known functions in cellular stress response have also been implicated in psychological stress-coping mechanisms, e.g., EGR1^14, 15^.

TOB has been proposed to regulate learning and memory, yet the mechanism is unknown^16, 17^. Notably, *Tob* is one of the early response genes after either neuronal depolarization in excitatory neurons or stress in humans^18, 19^. In addition, TOB protein expression is elevated in hippocampus and cerebellum after behavioral tests like fear conditioning and rotarod tests in rats, respectively^16, 17^. Moreover, decreased *Tob* gene expression has been correlated with depression^20^. Taken together, this suggests that TOB participates in neuronal molecular machinery and behavioral phenotypes. On the other hand, we previously showed that TOB exhibits a unique transient elevation after exposure to UV stress, halting apoptosis, and then eliciting an apoptotic signal after undergoing proteasome-dependent degradation^21^. In this manner, TOB allows cells to recover through DNA repair mechanisms^22^. Furthermore, overexpression of TOB in human bronchial epithelial cells leads to protection from ionizing radiation-mediated cell death, increased ERK phosphorylation, and induced expression of DNA repair proteins^23^. Stimulation using BMP-2, which induces oxidative stress, led to increased TOB protein expression^24, 25^. This suggests that TOB contributes to stress machinery, mostly protective, at both the cellular and molecular levels. However, TOB’s function in psychological stress remains enigmatic.

Utilizing *Tob*-KO mice, we show that TOB has a functional role in stress coping behavior in the brain by regulating hippocampal connectivity, neuronal excitability, and temporal molecular changes induced by stress. Increased TOB protein expression in mouse brain after exposure to acute stress, accompanied by the abnormal behavioral phenotype in *Tob*-KO mice, reveals TOB as key molecular effector in the brain’s stress resilience.

## Results

### TOB protein increases in response to stress

TOB’s function as an anti-proliferative protein is well known, but the potential role it plays in regulating brain function is not well understood^16–18, 20, 26, 27^. With this objective, we analyzed levels of TOB protein in mouse brain. We show that TOB is ubiquitously expressed across various regions of mouse brain (Fig. 1A). Effector proteins controlling neuronal functions usually show synaptic expression patterns^28^. Likewise, TOB protein is localized in the neuronal synaptic fraction, including synapto-neurosomes, pre-synapses and post-synapses (Fig. 1B). TOB responds to cellular stress, neuronal activation, and glucocorticoid stimulation^18, 19, 21^. Therefore, we examined whether TOB protein levels change in response to acute psychological stress. Restraint stress and inescapable electric shock are widely used models of acute psychological stress^29, 30^. The hippocampus is associated with responses to acute stress^31^ and in the hippocampus, TOB protein increased by 49.5% and 59.3% at 3 and 5 hours (F_4,15_=6.050, p=0.0042; No stress vs 3h p=0.0205 and No stress vs 5h p=0.0058) post-exposure to 30 min of restraint stress, respectively, compared to non-stressed mice (Fig. 1C). Additionally, hippocampal TOB increased by 63.8% (F_4,10_=5.849, p=0.0108; No stress vs 1hr p=0.0221) 1 h after mice were introduced to inescapable electric shock stress (Fig. 1D). ERK phosphorylation levels also increased after acute stress, concurrently with TOB expression. Thus, TOB is expressed in the mouse brain and its expression is increased following acute stress.

**Fig.1:**
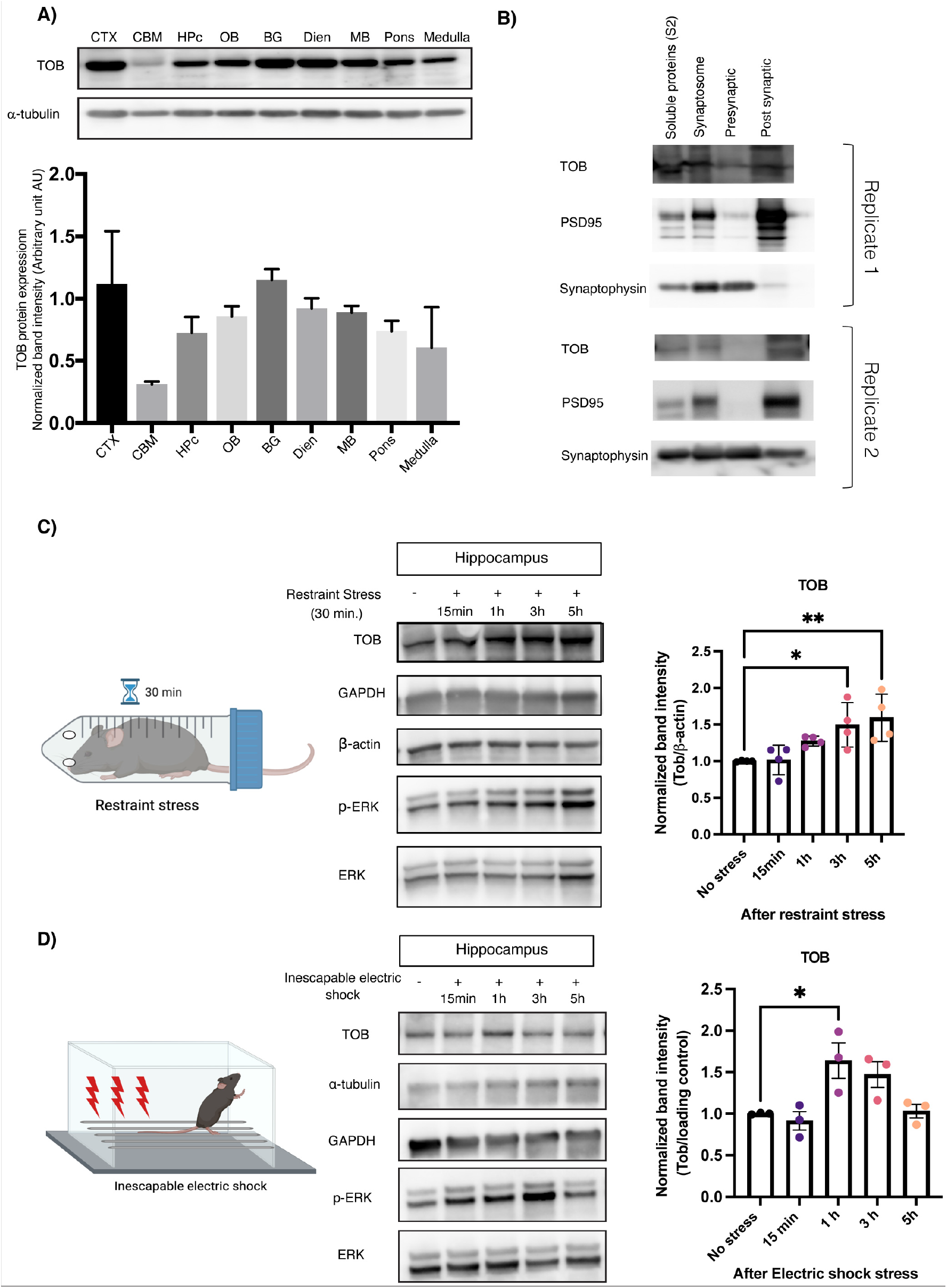
TOB protein expression levels increase in response to stress. A) Expression patterns of TOB in lysates of different mouse brain regions (n=3). B) Immunoblotting of TOB, synaptophysin, and PSD-95 in hippocampal fractionated lysates: soluble fraction S2, synaptoneurosomes, pre-synaptic, and post-synaptic fractions. C) Western blotting of TOB expression levels in hippocampal lysates without stress and after 30 min of restraint stress at different times: 15 min, 1 h, 3 h, 5 h after stress exposure (n=4). D) Western blotting of TOB expression levels in hippocampal lysates without stress and after inescapable electric shock for different durations: 15 min, 1 h, 3 h, 5 h post-exposure to stress (n=3). One-way analysis of variance (ANOVA) followed by Dunnett’s post-hoc correction for multiple comparisons: statistical significance *p<0.05 **p<0.01 when compared to control (No stress). Data are presented as means ±SEMs.

### Deletion of *Tob* alters the brain’s functional connectivity

Next, we sought to investigate the functional influence of *Tob* deletion on brain activity with resting-state functional magnetic resonance imaging (rs-fMRI) in the awake state with habituation to a small rodent MRI scanner. Awake resting-state fMRI for small animals allows us to observe brain-wide synchronization of hemodynamic signals across multiple brain regions. Prior to awake rs-fMRI sessions, *Tob*-KO and WT groups underwent habituation training for 7 days in a mock MRI environment with MRI scanning sounds (Fig. 2A-C). In order to check functional association with hippocampal CA1 and mPFC, we performed seed-based functional connectivity (FC) analysis and contrasted *Tob*-KO and WT groups. The seed-based FC analysis with bilateral CA1 revealed the statistical significance of FC with Dentate Gyrus (DG; p < 0.05 with FDR correction, Fig. 2D) and the primary somatosensory area (PSSA; p < 0.05 with FDR correction; Fig. S1). The previous analysis of CA1 revealed positively higher FC with DG and negatively higher FC with PSSA (p < 0.01 by Mann-Whitney U test; Fig. 2E). Furthermore, the seed-based FC analysis with mPFC revealed the statistical significance with DG (p < 0.05 with FDR correction; Fig. 2F) and SMA (SMA; p < 0.05 with FDR correction; Fig. S1). The previous analysis of mPFC also showed negatively higher FC with DG and SMA (p < 0.01 by Mann-Whitney U test; Fig. 2G). Our results imply that Tob KO may influence functional associations within hippocampal complex (HC) and between HC and mPFC.

**Fig.2:**
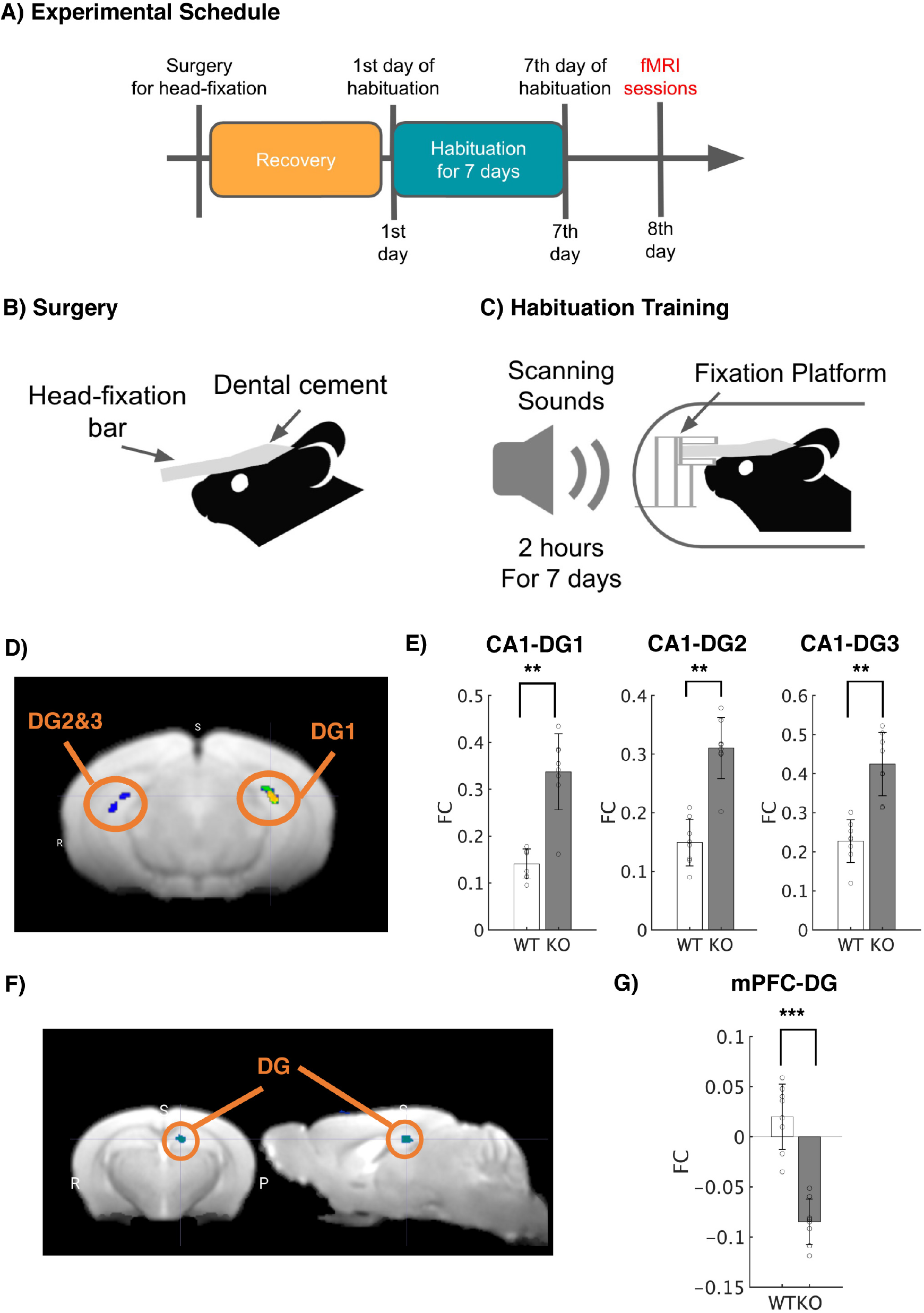
Deletion of *Tob* alters brain functional connectivity. A) Experimental Schedule. After surgery to introduce a head-fixation bar on the skull, mice were allowed to recovery. After recovery periods, mice underwent habituation training 2 h for 7 days prior to fMRI sessions. B) Surgery. A plastic head-fixation bar was mounted on the skull with dental cement. C) Habituation Training. In order to reduce scanning stress, mice were fixated with a fixation platform, and their bodies were constrained in a plastic tube. They were exposed to scanning sounds for 2 h for 7 days in order to reduce stress responses. D) Statistical functional map with the seed region, CA1. Average BOLD signals were extracted from bilateral CA1. Seed-based functional connectivity was performed, and a statistical map was visualized (p < 0.05 after cluster correction; Fig. S1). E) Functional connectivity with the bilateral CA1 seed. Seed-based FC analysis revealed statistically significant FC in CA1-DG1-3 in the Tob KO group with Mann–Whitney U test (** p < 0.01 with Bonferroni Correction). F) Statistical functional map with the seed region, mPFC. Average BOLD signals were extracted from the mPFC. Seed-based functional connectivity was performed, and a statistical map was visualized < 0.05 after cluster correction; Fig. S1). G) Functional connectivity with the mPFC seed. Seed-based FC analysis revealed statistically significant FC in mPFC-DG in the Tob KO group (*** p < 0.001 with Bonferroni Correction).

### Altered excitatory/inhibitory balance in *Tob*-KO hippocampal slices

To test whether *Tob* deletion alters synaptic function, we performed whole-cell patch clamp recordings in hippocampal CA1 pyramidal neurons in acute brain slices. We first investigated excitatory synaptic transmission in hippocampal CA1 pyramidal neurons. We found that *Tob* deletion significantly increased the amplitude (Fig. 3B), but not the frequency (Fig. 3C) of spontaneous miniature excitatory postsynaptic currents (mEPSC) when compared to WT mice. This selective change in mEPSC amplitude suggests that TOB deletion may enhance the number of postsynaptic receptors and/or the size of released presynaptic vesicles.

**Fig.3:**
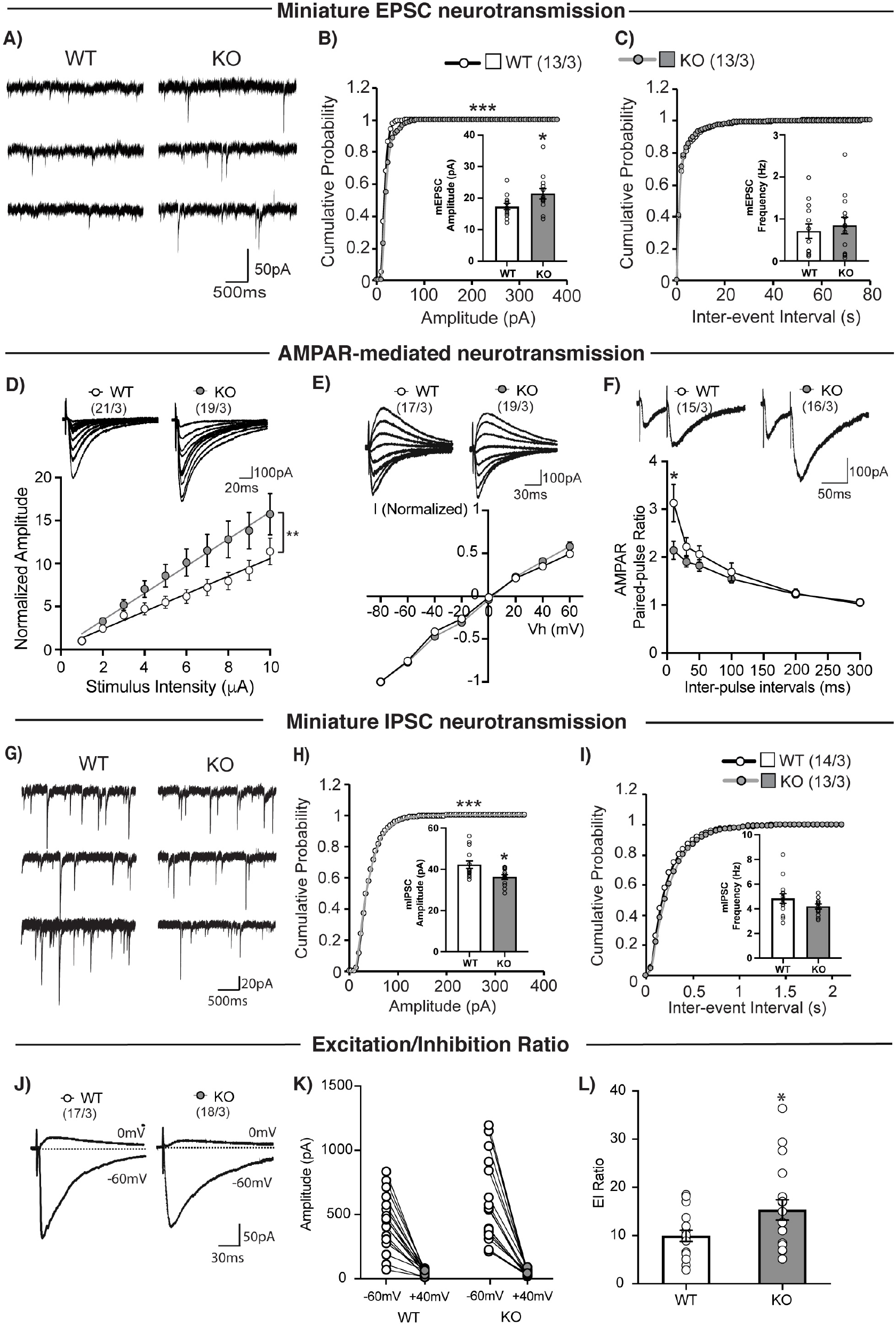
Altered excitatory/inhibitory balance in *Tob*-KO hippocampal slices. A) Representative traces of mEPSCs recorded from hippocampal pyramidal neurons of wild-type (WT, left traces) and *Tob* knockout (KO, right traces) at Vh of −70 mV in the presence of 1 μM tetrodotoxin and 100 μM PTX. Scale bars, 50 pA and 500 ms. B) Cumulative distribution plots and summary bar graphs for mEPSC amplitude (inset shows the average mEPSC amplitude) in CA1 hippocampal pyramidal neurons of wild-type (white column) and *Tob*-KO (grey column) mice. \*\*\**p* < 0.0001 by Kolmogorov-Smirnov test in the cumulative distribution plot and \**p* = 0.0453 by unpaired Student’s *t* test in the bar graph. C) Cumulative distribution plots and summary bar graphs for the mEPSC inter-event interval (inset shows the average of mEPSC frequency) in CA1 hippocampal pyramidal neurons of wild-type (white column) and *Tob*-KO (grey column) mice. *p* = 0.4611 by Kolmogorov-Smirnov test in the cumulative distribution plot and *p* = 0.6164 by unpaired Student’s *t* test in the bar graph. D) Sample traces (upper panel) and summary plots for the input-output relationship of AMPA receptor-mediated responses recorded from wild-type (open circles) and *Tob*-KO (grey circles) mice. Scale bars, 100 pA and 20 ms. \*\**p* = 0.0039 by Mann-Whitney U test. E) Sample traces (upper panel) and summary plots for the I-V curve of AMPA receptor-mediated responses recorded from wild-type (open circles) and *Tob*-KO (grey circles) mice. Scale bars, 100 pA and 30 ms. F) Sample traces with 50-ms inter-pulse interval (upper panel) and summary plots for paired-pulse ratio of AMPA receptor-mediated responses at 10, 30, 50, 100, 200 and 300 ms inter-pulse intervals recorded from wild-type (open circles) and *Tob*-KO (grey circles) mice. Scale bars, 100 pA and 50 ms. \**p* = 0.0139 by by Mann-Whitney U test; \*\*\**p* < 0.0011 by Two-way ANOVA with Sidak’s multiple comparisons test. G) Representative traces of mIPSCs recorded from hippocampal pyramidal neurons of wild-type (WT, left traces) and *Tob*-KO (KO, right traces) at Vh of −70 mV in the presence of 1 μM tetrodotoxin and 100 μM PTX. Scale bars, 20 pA and 500 ms. H) Cumulative distribution plots and summary bar graphs for mIPSC amplitude (inset shows the average of mIPSC amplitude) in CA1 hippocampal pyramidal neurons of wild-type (white column) and *Tob*-KO (grey column) mice. \*\*\**p* < 0.0001 by Kolmogorov-Smirnov test in the cumulative distribution plot and \**p*= 0.0120 by unpaired Student’s *t* test in the bar graph. I) Cumulative distribution plots and summary bar graphs for the mIPSC inter-event interval (inset shows the average of mIPSC frequency) in CA1 hippocampal pyramidal neurons of wild-type (white column) and *Tob*-KO (grey column) mice. *p* = 0.9311 by Kolmogorov-Smirnov test in the cumulative distribution plot and *p* = 0.1633 by unpaired Student’s *t* test in the bar graph. J) Representative traces of evoked EPSCs at Vh = −60 mV and evoked IPSCs at Vh = 0 mV in wild-type (left) and *Tob*-KO (right) mice. Scale bars, 50 pA and 30 ms. K) Amplitudes of evoked EPSCs at Vh of −60 mV and evoked IPSCs at Vh of 0 mV at each of individual recorded WT and *Tob*-KO hippocampal pyramidal neurons. L) Average excitation/inhibition ratio from WT (open column) and *Tob*-KO (grey column). \**p* = 0.0343 by unpaired Student’s *t* test. Data are expressed as means ±SEMs. Total numbers of cells recorded / total numbers of mice used are indicated in parentheses.

We further explored whether AMPA and NMDA receptors, the two major glutamate receptor classes mediating fast excitatory synaptic transmission, contribute to the dysregulation after TOB deletion (Fig. 3A-C). We characterized AMPA receptor-mediated synaptic transmission at CA3-CA1 synapses in the hippocampus by assessing the input (stimulation intensity)-output (EPSC amplitude) efficiency and voltage dependence of synaptic AMPA-mediated synaptic responses in the presence of the NMDA receptor antagonist, D-APV. Slopes of the linear fit for individual AMPA-mediated input-output experiments were significantly different in KO mice compared to those of WT (p = 0.0039) (Fig. 3D). No apparent difference between genotypes was found in the current-voltage (I-V) curve (Fig. 3E). The rectification index of AMPA receptor-mediated responses from *Tob*-KO was also comparable to that of WT (WT: 0.861 ± 0.08; KO: 0.783 ± 0.07; p = 0.568 with the Mann-Whitney U test). Moreover, AMPA receptor-mediated, paired-pulse facilitation was slightly increased in KO synapses (Fig. 3F), most pronouncedly at 10-ms inter-pulse intervals (p = 0.032). This suggested that TOB deletion resulted in an increase in the number of mature postsynaptic AMPA receptors without changing the AMPA receptor subunit composition.

We next investigated NMDA receptor-mediated synaptic transmission at CA3-CA1 synapses in the hippocampus. We measured the input-output relationship and I-V curve of synaptic NMDA receptor-mediated synaptic responses in the presence of the AMPA receptor antagonist, NBQX. We found no apparent differences between genotypes (Fig. S2 A-B). The rectification index of NMDA receptor-mediated responses from *Tob*-KO was also similar to that of wild-type (WT: 2.959 ± 0.444; KO: 2.032 ± 0.136; p = 0.135 with the Mann-Whitney U test).

We further examined inhibitory synaptic functions in hippocampal CA1 pyramidal neurons. The amplitude (Fig. 3H), but not the frequency (Fig. 3I) of miniature inhibitory postsynaptic currents (mIPSCs) were reduced in *Tob*-KO mice, compared to that of wild-type mice (Fig. 3G-I). These results indicate that TOB deletion affects both excitatory and inhibitory synaptic transmission.

Next, we directly estimated the ratio of excitation to inhibition in hippocampal pyramidal neurons. We first recorded AMPA receptor-mediated EPSCs at a holding potential of −60 mV, equivalent to the Cl^-^ equilibrium potential. Then we recorded GABA_A_ receptor-mediated IPSCs at a holding potential of 0 mV, which is the reverse potential of AMPA and NMDA receptors. We then calculated the ratio of EPSC amplitude to that of IPSC amplitude (E/I ratio) and found that the E/I ratio was markedly increased in *Tob*-KO mice (Fig. 3J-L).

### *Tob*-KO mice show abnormal stress-related behavior

Contextual fear conditioning includes exposing mice to aversive acute stress caused by inescapable electric shocks. Then brain regions respond by associating the context to such a stimulus. Fearful mice freeze when exposed to the same context in which conditioning occurred. On the other hand, contextual fear extinction is the subsidence of fear response due to repetitive exposure to the same context without shock presentation^32^. After fear conditioning, *Tob*-KO mice exhibited increased contextual fear freezing (F_3,30_=10.77, p<0.0001 for genotype effect; F_15,150_=2.727, p=0.0010 for time x genotype effect; WT vs KO at Day 1 p<0.0001, Day 2 p<0.0001, Day 3 p=0.0003, Day 4 p=0.0342) (Fig. 4A). Overexpression of TOB in the hippocampus rescued the abnormal fear phenotype as *Tob*-KO (AAV_mTob) did not exhibit significant fear compared to *Tob*-WT(AAV_mTob) any time after conditioning. Additionally, KO mice rescued through overexpression of AAV_mTob, showed significantly less freezing when compared to KO (KO(AAV_mTob) vs KO at Day 1 p<0.0001, Day 2 p<0.0001, Day 3 p=0.0058) (Fig. 4A, S3 A-D). TOB deletion in the hippocampus causes an increased fear response to an aversive context and decreased extinction, which was reversed by re-expression of TOB.

**Fig.4:**
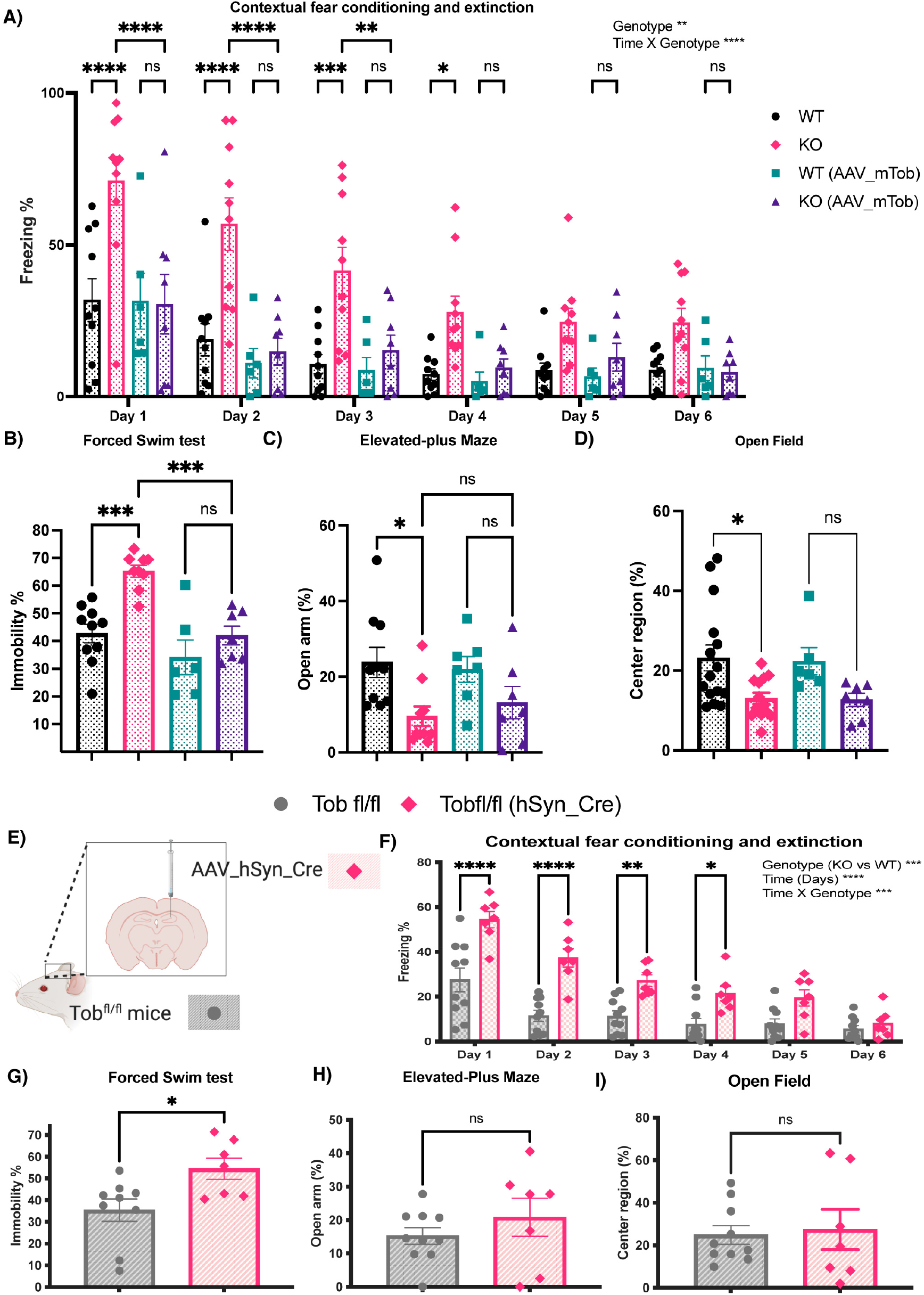
*Tob*-KO mice show hippocampal-mediated abnormal stress-related behavior. Behavioral analyses in *Tob*-WT and KO mice and after overexpression of mouse TOB using AAV (hSyn-mTob) (A-D). A) Contextual fear conditioning and extinction expressed as percentage of time spent freezing. Two-way ANOVA followed by Bonferoni’s *post-hoc* test for multiple comparisons. B) The forced swim test presented as a percentage of immobile time. One-way ANOVA followed by Bonferoni’s *post-hoc* test for multiple comparisons. C) Elevated-plus maze showing the percentage of time spent in open arm. One-way ANOVA followed by Bonferoni’s *post-hoc* test for multiple comparisons. D) Open field test showing the percentage of time spent in center region. One-way ANOVA followed by Bonferoni’s *post-hoc* test for multiple comparisons. Behavioral analyses in hippocampal-specific *Tob*-KO mice (E-I). E) Schematic diagram showing the method for generation of hippocampal-specific *Tob*-KO (hsTobKO) mice through injection of adeno-associated virus expressing Cre recombinase under the hSyn promoter (AAV_hSyn_Cre) in mice having LoxP sequences flanking both sides of the *Tob* gene (*Tob*^fl/fl^). F) Contextual fear conditioning and extinction in hsTobKO presented as percentage of time showing freezing. Two-way ANOVA followed by Bonferoni’s *post-hoc* test for multiple comparisons. G) The forced swim test is presented as percentage of time spent immobile. H) The elevated-plus maze showed as the time spent in the open arm. I) Open field test showing the percentage of time spent in the center region. Unpaired t-test. All values represent means ± SEMs. ns non-significant, * p<0.05, **p<0.01, ***p<0.001, ****p<0.0001

The forced swim test, in which immobility is associated with increased despair, is widely used to test depression-like behavior, but it is also an efficient test of the ability to cope with stress^33^. *Tob*-KO mice showed increased immobility in the forced swim test (F_3,28_=13.50, p<0.0001; WT vs KO p=0.0003). Re-expression of TOB in the hippocampus of *Tob*-KO mice reduced immobility (KO(AAV_mTob) vs KO p=0.0008; WT(AAV_mTob) vs KO(AAV_mTob) p>0.9999) (Fig. 4B). Similarly, we observed increased immobility by *Tob*-KO mice in the tail suspension test, which was rescued by TOB overexpression in the hippocampus (Fig. S3E). This shows that TOB in the hippocampus is important for coping with stress.

Since anxiety is usually observed in models showing abnormal stress coping mechanisms, we next analyzed anxiety in our mouse model. *Tob*-KO mice spent less time in the open arm of the elevated-plus maze, an indication of increased anxiety (F_3,30_=3.948, p= 0.0174; WT vs KO p=0.0283) (Fig. 4C). Unlike in fear conditioning, TOB re-expression in hippocampus did not decrease anxiety in KO mice, as time spent in the open arm was not significantly different.

In the open-field test, *Tob*-KO mice spent less time in the center region than WT mice (F_3,37_=4.263, p=0.0111; WT vs KO p=0.0309; WT(AAV_mTob) vs KO(AAV_mTob) p=0.3621) (Fig. 4D, S3 F-H). Although the time spent in center region was still low after overexpression of AAV_mTob in the hippocampi of KO mice, there was no significant difference between WT(AAV_mTob) and KO(AAV_mTob). Therefore, we believe that the increased anxiety in Tob KO mice is not hippocampus-dependent.

In order to identify specific brain areas associated with *Tob* behavioral deficiencies, we generated hippocampus-specific *Tob*-KO mice (hsTobKO) using the Cre-loxP system. First, *loxP* sequences flanking exon2 were inserted in the *Tob* gene (*Tob^fl/fl^*) (Fig. S4A). Adeno-associated virus expressing Cre recombinase under the human synapsin 1 (*hSyn*) promoter (AAV_hSyn_Cre) was injected into the dorsal hippocampus of *Tob^fl/fl^* mice to delete *Tob* specifically in this region in adult mice (Fig. 4E, S4B-C). We then analyzed behavior of hsTobKO mice. Freezing in the same context, where mice had undergone fear conditioning, and subsequent extinction trials were increased after TOB deletion in hippocampus (F_1,14_=26.11, p=0.0002 for genotype effect; F_5,70_=4.701, p=0.0009 for time x genotype effect; AAV_hSyn_Cre vs Tob^fl/fl^ at Day 1 p<0.0001, Day 2 p<0.0001, Day 3 p=0.0148, Day 4 p=0.1301) (Fig. 4F). Additionally, hsTobKO did not show abnormal cued fear (Fig. S4D-E). Depression-like behavior was observed in hsTobKO mice in that immobility time was higher during the forced swim test (t-test p=0.0193; Fig. 4G), and tail suspension test (Fig. S4F). On the other hand, anxiety levels were normal, and time spent in the open arm did not differ in hsTobKO and *Tob*^fl/fl^ mice (t-test p=0.3329) (Fig. 4H). Also, no abnormal anxiety was observed in hsTobKO mice, as there was no change in time spent in the center of the open field test (t-test p=0.0972) was observed (Fig. 4I, S4G-H). These results show that TOB in the hippocampus is important for normal fear and depression behaviors.

### Abnormal transient transcriptional profile in hippocampus of Tob KO mice and suppressed stress-induced LCN2 expression induced after fear conditioning

Stress stimulates neuronal activation, which in return induces changes in the underlying molecular signaling pathways. Abnormal responses at the molecular level would lead to aberrant stress coping behavior. Since *Tob*-KO mice showed increased contextual fear and abnormal extinction, we analyzed hippocampal transcriptomic changes after fear conditioning. To analyze rapid changes resulting from aversive stimuli in fear conditioning, we performed RNA sequencing on hippocampal RNA from mice culled at 15 min, 1 h, and 3 h post-conditioning, in addition to naïve mice. When compared to WT mice, differentially expressed genes in *Tob*-KO mice were 3,3,2 upregulated and 2,1,3 downregulated genes for naïve mice, and 1 h and 3 h post-training, respectively (Fig. 5A). Strikingly, the greatest differential transcriptome changes in *Tob*-KO occurred 15 min after conditioning, in that 26 genes were upregulated and 11 genes were downregulated (EdgeR WT vs KO 15 min after fear conditioning, p<0.05, FDR<0.05). Therefore, TOB deletion altered the rapid change in the hippocampal transcriptome after fear conditioning.

**Fig.5:**
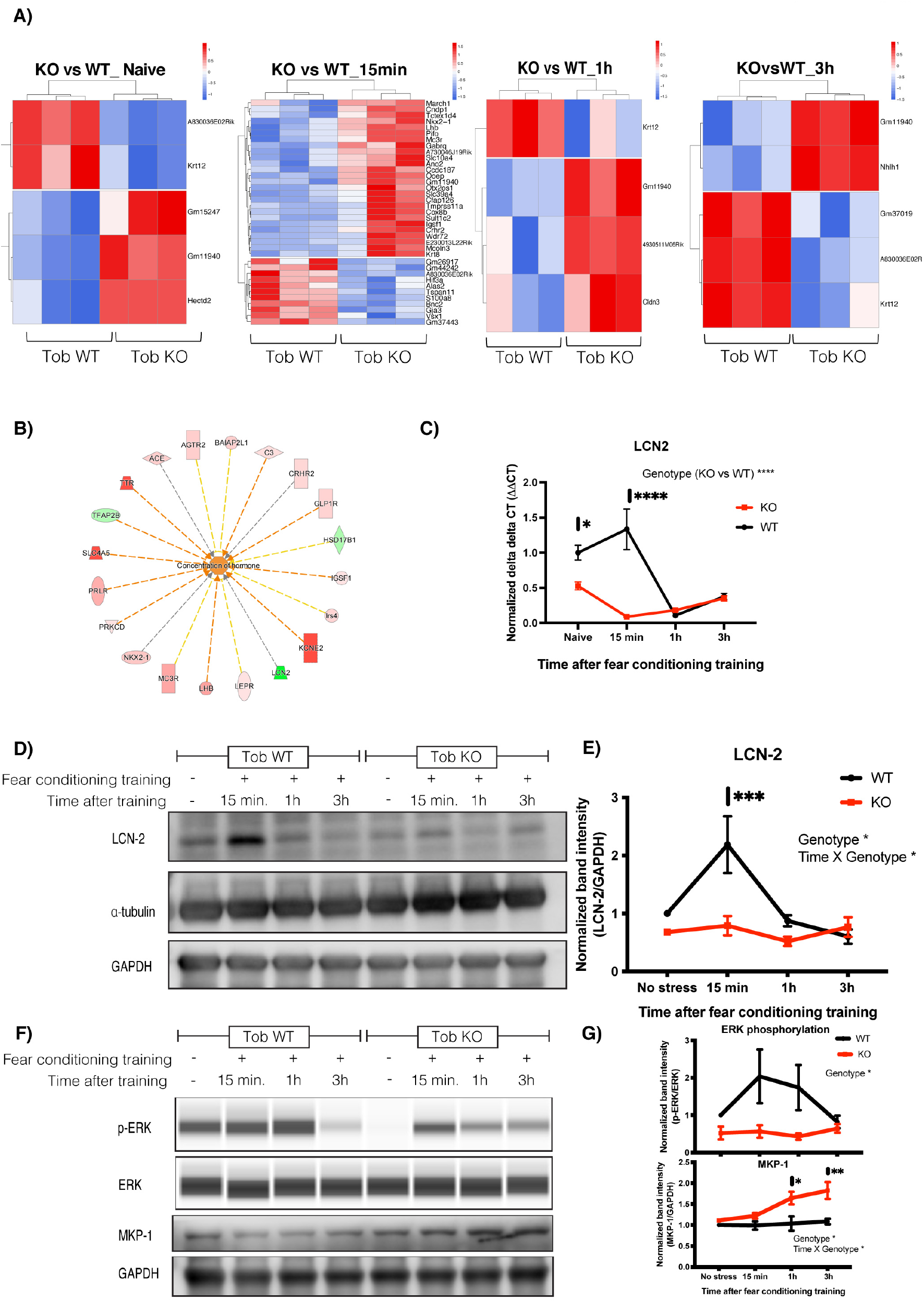
Abnormal transient transcriptional profile in hippocampus of *Tob*-KO mice and suppressed stress-induced LCN2 expression induced after fear conditioning training. A) Heatmaps for differentially expressed genes in hippocampi of *Tob*-KO compared to *Tob*-WT mice using RNA sequencing without fear conditioning training (naïve) and 15 min, 1 h, 3 h after fear conditioning training (represented as z-scores of log raw counts, FC_upregulated_ >2 FC_downregulated_ <0.5, p<0.05, FDR<0.05). B) Pathway analysis for RNA sequencing candidates using IPA software showing activation of hormonal concentration in hippocampus of Tob KO mice at 15 minutes post-conditioning. C) Real-time PCR for lipocalin-2 (*Lcn2*) mRNA in hippocampus of *Tob*-WT and KO naïve mice and 15 min, 1 h and 3 h after fear conditioning. Two-way ANOVA followed by Bonferoni’s *post-hoc* test for multiple comparisons. D) Western blotting showing protein expression of LCN-2 in hippocampi of naïve *Tob*-KO mice and at 15 min, 1 h and 3 h after fear conditioning training. E) Normalized band intensity for LCN2 protein immunoblots. Two-way ANOVA followed by Bonferoni’s *post-hoc* test for multiple comparisons. F) Western blotting showing abnormal protein expression in hippocampi of mice lacking *Tob* before and after fear conditioning training at 15 min, 1 h and 3 h. G) Western blot band intensity quantification plots at different time points post-training compared to naïve *Tob*-WT (n=3). All values represent means ± SEMs. ns non-significant, * p<0.05, **p<0.01, ***p<0.001, ****p<0.0001

To understand the possible molecular pathway governing this transcriptomic change, we performed pathway analysis. Pathway analysis for differentially expressed genes 15 min after fear conditioning suggested upstream activation of hormone concentration, estrogen receptor 1 (*Esr1*) and dexamethasone-induced pathways with genes for receptors controlling the HPA axis and corticoids like *Mc3r, Crhr2, Avpr1a* and neuronal inflammation genes like *Lcn-2* (Fig. 5B, Fig. S5A-E).

Lipocalin-2 (*Lcn2*) was one of the transcripts downregulated in hippocampi of *Tob*-KO mice. To confirm this, we performed qRT-PCR, which showed lower mRNA levels in hippocampi of *Tob*-KO mice for naïve and 15 min post-fear conditioning (F_1,16_=27.4, p<0.0001 for Genotype effect; F_3,16_=14.22, p<0.0001 for Genotype x time effect; WT vs KO naïve p=0.0363, 15 minutes p<0.0001) (Fig. 5C). Lower *Lcn2* mRNA levels coincided with lower protein levels, in which fear conditioning-induced LCN2 protein expression after 15 min was suppressed in *Tob*-KO mice (F_1,4_=11.67, p=0.0269 for Genotype effect; F_3,12_=5.199, p=0.0157 for Time x Genotype; WT vs KO 15 minutes p=0.0007) (Fig. 5D, 5E). These results show that TOB contributes to activation of stress-induced LCN2 transcription and subsequent translation.

Additionally, fear-induced ERK phosphorylation was inhibited in hippocampi of KO mice after fear conditioning (F_1,4_=12.6, p=0.0238 for Genotype effect) (Fig. 5F, 5G). On the other hand, MKP-1 protein levels were elevated in KO mice (F_1,3_=12.25, p=0.0395 for Genotype effect; F_3,9_=5.358, p=0.0216 for Time x Genotype; WT vs KO 1 h p=0.0124, 3 h p=0.0029). Accordingly, TOB deletion repressed stress-induced ERK phosphorylation and simultaneously increased MKP-1 protein levels.

## Discussion

Stress vulnerability and resilience differ among individuals, which suggests a strong influence of genetic factors. However, molecular mechanisms predisposing some individuals to stress-induced psychiatric disorders are not well explored. Here, we introduce TOB as a psychological stress-responsive gene that governs synchrony between brain regions that process emotional regulation. Additionally, *Tob*-deficient mice showed abnormal functional dynamics in the hippocampus and between the hippocampus and mPFC. Moreover, *Tob* deletion led to impaired hippocampal excitatory-inhibitory balance, accompanied by increased AMPAR- and decreased GABAR-mediated synaptic transmission. Additionally, TOB-deficient mice showed abnormal hippocampal-dependent contextual fear conditioning, extinction, and depression-like behaviors. *Tob* deletion resulted in abnormal transient hippocampal transcriptome profiling after fear conditioning in *Tob*-KO mice. Moreover, fear conditioning-induced *Lcn2* expression was suppressed in *Tob*-KO mice, mostly through the ERK-mediated pathway.

During its initial characterization, TOB was proposed to exhibit a transient response to stimulation^34^. Interestingly, *Tob* was recently classified as one of the early response genes^18, 19^, possibly due to the presence of three ATTTA motifs in the 3’UTR of the *Tob* gene, which is commonly observed in immediate early genes^35^. Our results show that hippocampal TOB levels rapidly increased after acute psychological stress. A similar pattern was observed in genes involved in the stress machinery pathway, like PAI-1 and tPA^36, 37^.

Our data show decreased hippocampal-medial-prefrontal functional connectivity in KO mice. Altered PFC connectivity to other brain regions predisposes depression, while enhancement is used to evaluate treatment efficiency^38^. Additionally, the mPFC has an inhibitory function in regulating emotional behavior, like reducing fear responses, extinguishing aversive memories, and suppressing the hypothalamic-pituitary adrenal (HPA) axis^39, 40^. Therefore, decreased functional connectivity in *Tob*-KO mice suggests a loss of inhibitory emotional control of the mPFC over the hippocampus. An aberrant increase of hippocampal CA1-DG functional connectivity in *Tob*-KO mice leads to enhanced synchronization and processing in the hippocampus. Together with the significant increase in the excitatory-inhibitory ratio, our data suggest that hippocampal activity is disproportionally increased. This is consistent with reports of decreased prefrontal and increased hippocampal activity in PTSD and depression patients^41–44^. One limitation to our fMRI analysis is that the amygdala, a central hub regulating emotional behavior, could not be included due to high noise originating from nearby veins and the air-filled ear canal^45, 46^. There are reports correlating fMRI with neuronal activity^47^. However, it was challenging to distinguish between excitatory and inhibitory neuronal activity in our study. Nonetheless, the change in hippocampal functional connectivity in *Tob*-KO mice suggests a causative change in neuronal activity. The mPFC is anatomically connected to the hippocampus through CA1^48^. Therefore, we decided to measure CA1 neuronal activity, as we speculated that it helps to regulate hippocampal and mPFC-hippocampal connectivity.

Our results show that TOB is expressed in the synaptic fraction, suggesting its possible function in synaptic neurotransmission. This is compatible with a single-cell sequencing study by Qiu et al. (2020)^18^, which showed a rapid increase in *Tob* expression in excitatory neurons after neuronal stimulation. Our results from *Tob*-deficient hippocampal slices show increased AMPAR-mediated and decreased GABAR-mediated neuronal transmission. Interestingly, increased expression of AMPARs and subsequent neuronal activity were observed in response to stress^49^. On the other hand, decreased inhibitory neurotransmission is a well-established etiology for depression, anxiety, and PTSD^50–52^. Also, inhibiting GABAergic neurons in CA1 altered the hippocampal-mPFC neuronal firing synchronization^53^. As expected, TOB-deficient slices show an elevated excitatory/inhibitory (E/I) ratio. A similar increase in E/I ratio is observed in stress and depression mouse models^54–56^. Taken together, the observed change in CA1 neuronal activity of *Tob*-KO mice is strongly linked to the altered functional connectivity between the hippocampus and prefrontal cortex. Additionally, the altered excitation and inhibition balance might be a consequence of decreased inhibitory synaptic transmission^50^. One limitation of our study is that fMRI imaging and electrophysiological recording were done on naïve non-stressed mice to analyze basal levels. This was necessary to evaluate abnormal factors leading to altered stress responses. Future measurement of neuronal activity at cellular and circuit levels after stress would offer more insight on the function of TOB in the brain’s dynamic stress network level.

Fear conditioning and extinction have been introduced as a model of PTSD, to assess emotional behavior in response to aversive stimuli^57^. A stress-induced increase in AMPARs was reported to enhance fear memory^58, 59^. Therefore, elevated hippocampal neurotransmission can be linked to the increased contextual fear conditioning and extinction of *Tob*-KO mice. Our rescue and hippocampal-specific knockout experiments demonstrate that enhanced fear in *Tob*-KO mice is hippocampal-dependent. Forced swim has been implemented to evaluate the ability to cope with inescapable stress^60, 61^. *Tob*-KO mice exhibit depression-like behavior when exposed to forced swim, this suggests that TOB may function in efficiently coping with stressors. Such depression-like behavior is consistent with previously reported low *TOB* mRNA levels in major depressive disorder (MDD) patients^20^. On the other hand, hippocampus-specific *Tob*-KO did not induce anxiety, and TOB re-expression in the hippocampus did not reduce it. Therefore, the increased anxiety in *Tob*-KO mice could be mediated by a brain region other than the hippocampus. This accords with studies showing that anxiety is mainly regulated by the amygdala and prefrontal cortex^62^. Taken together, TOB is important for intact hippocampal-mediated stress coping behaviors, namely fear and depression.

How does TOB execute stress-induced molecular functions in the hippocampus? TOB regulates gene transcription^63, 64^, this likely explains the increase in differentially expressed genes 15 min after fear conditioning in KO mice when compared to WT. The abnormal changes in hormone receptor levels of KO mice after fear conditioning suggest the activation of stress-induced hormonal pathway. *Ttr, Crhr2, Avpr1a* and *Mc3r* are among the activated genes that could be involved in this pathway. *Ttr* gene expression is regulated by glucocorticoids and estradiol ^65, 66^. *Crhr2* is activated by Urocotins I, II (stresscopin-related peptide), and III (stresscopin), which belong to the corticotrophin-releasing factor (CRF)-related peptides^67^. *Crhr2* expression in the brain has been correlated with fear^68^. Similarly, elevated Crhr2 was attributed to an inability to cope with stress, consequently predisposing the individual to PTSD or suicide^69, 70^. Interestingly, stimulating Crhr2 with different concentrations of agonist, led to either activation or inhibition of the HPA axis^71^. *Avpr1a* is another interesting candidate. Its activation leads to increased fear response and anxiety^72^. Additionally, AVPR1A contributes to activation of the HPA-axis by increasing ACTH and corticosterone levels^73^. Lastly, Melanocortin 3 receptor (*Mc3r*) is another candidate activated by melanocortin peptides, namely alpha, beta, and gamma-melanocyte-stimulating hormones (MSHs) and adrenocorticotrophic hormone (ACTH))^74^. MC3R activation has been linked to increased anxiety and stress response^75^. Collectively, the increased mRNA expression of hormone receptor levels in hippocampi of *Tob*-KO mice after fear conditioning may give rise to hormone-mediated abnormal behavior, partly through the hyperactivated HPA axis. This might also be induced by decreased inhibitory neurotransmission, which increases HPA axis activity^76^, anxiety, and responses to stress^77^.

Fear conditioning induced a transient increase in LCN2 expression in hippocampi of WT mice, which was inhibited by *Tob* deletion. Like *Tob*, *Lcn2* is an acute phase gene^78^ that shows increased hippocampal expression in response to restraint stress^79^. Additionally, LCN2 deletion in mice caused anxious- and depressive-like behaviors^80^, which resemble those of *Tob*-KO mice. Lower LCN2 expression in TOB-deficient hippocampus can be linked to the observed increase in CA1 excitatory neurotransmission, as LCN2 deficiency induces hippocampal neuronal excitability^79^. Therefore, TOB functions in response to stress may be partially mediated through LCN2. However, until now there has been no evidence showing any interaction between TOB and LCN2. ERK phosphorylation might be the missing link between TOB and LCN2. We observed decreased stress-induced ERK phosphorylation in hippocampus of *Tob*-KO mice. Decreased ERK phosphorylation has been attributed to stress-induced depression^81^. The interaction between TOB and ERK is bidirectional, in that ERK phosphorylates TOB and TOB impacts ERK phosphorylation^23, 82, 83^. On the other hand, ERK phosphorylation induces LCN2 expression^84^. Therefore, it is highly suggestive that decreased LCN2 levels after stress in *Tob*-KO mice are due to altered ERK phosphorylation. One of the upstream phosphatases controlling ERK phosphorylation is MKP-1, which is overexpressed in hippocampi of *Tob*-KO mice. This is consistent with previous reports that MKP-1 expression is induced by cellular stress and acute glucocorticoid treatment^85^, to inactivate MAP kinases such as ERK^86^.

In summary, this study demonstrates increased TOB levels after acute stress and highlights its function in the hippocampus by maintaining normal fear and depressive behaviors. We also show that TOB regulates the rapid transcriptional response after acute stress, hippocampal connectivity, and synaptic transmission. These observations set the stage for future use of TOB as a stress biomarker or vulnerability predictor for individuals prone to stress.

## Materials and Methods

### Animals

Mice with a C57BL/6J genetic background were used in this study. *Tob*-KO mouse generation and validation were described by Yoshida et al., 2000^64^. Floxed *Tob* mice (Accession No. CDB0044E) were generated by insertion of LoxP sequences spanning exon2 of the *Tob* gene with detailed procedure described in supplementary information. Genotyping to detect insertion at the 5’ end employed primers: FW 5’-TGAGAGCCCTTGGCATGG −3’ REV 5’-ATACCACTTCCCAGCAGG −3’and at 3’ end using: FW 5’-GGAATAATGGAAGGCAGG −3’ REV 5’- CCTCCTATCACCTGGCTC −3’. Mice with homozygous LoxP insertions (Floxed *Tob* mice, Tob^fl/fl^) were used for experiments after backcrossing with C57BL/6J mice for at least 5 generations. All mice were housed under controlled temperature and a 12-h light/dark cycle. All animal experiments were performed following guidelines for experimental animals and approved by the Animal Care and Use Committee, Okinawa Institute and Science Technology Graduate University (OIST), Japan.

### Restraint Stress

Mice were restrained in 50-mL Falcon centrifuge tubes with conical bottoms (Corning, USA) for 30 min. Holes were drilled in the tubes to allow respiration, while tube caps had one hole to let their tails pass through. After restraint stress, mice were returned to their home cages for indicated times, to be sacrificed for collection of hippocampi for protein extraction.

### Inescapable electric shock

Mice were exposed inescapable electric shocks as described in the training procedure for “Fear Conditioning and Extinction”, and then returned to their home cages until sacrifice and collection of hippocampi at the indicated times.

### Western blotting

Hippocampal tissues were lysed using ice-cold lysis buffer containing 0.3% SDS, 1.67% Triton X-100, 50 mM Tris-HCl pH7.4, 150 mM NaCl, 1 mM EDTA, 1 mM EGTA, 10% glycerol, Halt Protease inhibitor cocktail (ThermoFisher, USA), and phosphatase inhibitor cocktail PhosSTOP (Roche, Switzerland). Synaptic fractionation was done following a detailed published protocol^87^. Electrophoresis was performed using 7.5 or 12% TGX Acrylamide gels (Bio-rad, USA) following standard protocols. Proteins were transferred to PVDF membranes using Trans-Blot Turbo Transfer (Bio-Rad, USA) and blocked in TBS buffer containing 5% BSA and 0.1% Tween-20. Antibodies were diluted in Can Get Signal immunoreaction enhancer solution (Toyobo, Japan) and incubated according to the manufacturer’s protocol. Antibodies used were anti-Tob mouse monoclonal antibody as described by Matsuda et al., 1996, anti-Tob rabbit polyclonal antibody (RpAb) (Sigma-Adrich, USA), anti-ERK1/2 rabbit monoclonal antibody, anti-(p-ERK1/2) RpAb, anti-Cre RpAb (Cell Signaling, USA), anti-MKP-1 RpAb (Santa Cruz, USA), anti-LCN2 goat polyclonal antibody (R&D systems, USA). Chemiluminescent signals were generated using Immobilon (Millipore, USA) and detected using ImageQuant LAS4000 (GE healthcare, USA). For reprobing, Restore Plus stripping buffer (ThermoFisher, USA) was used. Band intensities were quantified using Image Studio Lite software (Li-Cor, USA). Automated Simple Wes system was used to quantify ERK phosphorylation levels, according to the manufacturer’s instructions using the 12–230 kDa separation module and anti-rabbit detection module (ProteinSimple, USA).

### Functional magnetic resonance imaging (fMRI)

Detailed head-fixation bar mounting surgery and MRI imaging procedures are described in the supplementary information.

### Functional connectivity analysis

The pre-processed and denoised time series data were used for a seed-based FC analysis with CONN17. Regions of interest (ROIs) including CA1, DG and mPFC were chosen. Seed-based functional connectivity (FC) analysis was performed to compare FC between the Tob-KO group and the control group. Seed-based FC analysis was composed of two steps. First, Pearsons’ correlation between a time series of an average seed ROI and each voxel in images was calculated, and regional clusters were formed by thresholding statistical significance (uncorrected p-value < 0.001) between two groups. In the second step, formed clusters were further statistically corrected with a positive false discovery rate (pFDR; p < 0.05).

### Electrophysiological recording

Electrophysiological recordings were performed as described by Etherton et al.^88^ and are detailed in the Supplementary materials and methods.

### Quantitative real-time PCR

Total RNA was extracted from mouse hippocampi using Isogen II (Nippon Gene, Japan) following the manufacturer’s protocol. Reverse transcription was performed using PrimeScript II 1st strand cDNA Synthesis Kit (Takara, Japan) following the manufacturer’s protocol. Real-time PCR was performed using TB Green Premix Ex Taq II (Takara, Japan) and ViiA7 Real-Time PCR system (Applied Biosystems, USA). Relative mRNA expression was determined by the ΔΔCT method and *Gapdh* mRNA levels were used for normalization. Primers used were: *Gapdh* FWD 5’- ctgcaccaccaactgcttag −3’ REV 5’- gtcttctgggtggcagtgat −3’; *Lcn2* FWD 5’- ccccatctctgctcactgtc −3’ REV 5’ - tttttctggaccgcattg −3’; *Crhr2* FWD 5’- aagctggtgatttggtggac - 3’ REV 5’-ggtggatgctcgtaacttcg −3’; *Avpr1a* FWD 5’- gctggcggtgattttcgtg −3’ REV 5’- gcaaacacctgcaagtgct −3’; Mc3r FWD 5’- tccgatgctgcctaacctct −3’ REV 5’- ggatgttttccatcagactgacg −3’; *Ttr* FWD 5’- agccctttgcctctgggaaga −3’ REV 5’- tgcgatggtgtagtggcgatgg −3’.

### RNA sequencing

Intact poly(A) RNA was purified from 1 μg of total RNA using an NEBNext^®^ Poly(A) mRNA Magnetic Isolation Module (New England Biolabs, USA) and following the manufacturer’s protocol. Library preparation was performed using NEBNext® Ultra II Directional RNA Library Prep Kits for Illumina (New England Biolabs, USA), according to the manufacturer’s protocol with 8 PCR cycles. Library sizes were checked using microfluidic-based electrophoresis LabChip GX Touch (Perkin Elmer, USA) and concentrations were checked using Qubit 1X dsDNA HS (ThermoFisher, USA) and then pooled after concentration adjustment. 150-bp paired-end RNA sequencing was performed using a NovaSeq 6000 SP flow cell (Illumina, USA).

Analysis was done using fastq files containing paired-end sequencing reads and analyzed using nf-core/rnaseq pipeline v2.0^89^, which were mapped to the GRCm38 genome database using STAR aligner (v2.6.1d)^90^. Mapped genes were then further analyzed using OmicsBox software (v1.4.11) for counting using HTSeq (v0.9.0)^91^ and differential gene expression analysis using the package EdgeR (v3.11)^92^. Reads were normalized using the Trimmed Mean of M-values (TMM) normalization method and a cut-off of at least 0.2 counts per million (CPM) in two samples was selected. Differentially expressed genes (DEGs) were statistically tested using EdgeR’s exact test, and genes with FDR≤0.05, p-value≤0.05 and fold change (FC) ≥2 or ≥-2 were used for further analysis. Pathway analysis was performed for genes 2-fold up- or down-regulated with p-value < 0.05 using Ingenuity Pathway Analysis (IPA) software (Qiagen, USA).

Raw and pre-processed transcriptomic data files described in the current study are publicly available in NCBI GEO under accession number GSE186101.

### Behavior

Behavioral analyses were performed using male mice 8-12 weeks old. All experiments were performed by experimenters blinded to genotype during testing. All software for analysis was from O’Hara & Co Ltd., which has been modified in the public domain (National Institutes of Health (NIH) Image J program).

### Fear Conditioning and extinction

Fear conditioning and extinction were performed as described previously by Pibiri et al., 2008^93^ with minor modifications. Briefly, on training day, mice were placed in a conditioning chamber (CL-3002L, O’Hara & Co Ltd., Japan) for 2 min to habituate, and then presented with a conditioning stimulus (CS) of a 65-dB tone for 30 s, co-terminated with an unconditioned stimulus (US) of 0.5 mA, a 2-s foot shock. The tone and foot shock were repeated 3 times at 2-min intervals. Mice were returned to their home cages 30 s after the last shock. During the contextual fear test (Day 1), mice were placed in the chamber for 5 min without any tone or shock presentation. During cued fear conditioning, mice were placed in a novel chamber for 6 min and allowed to explore then presented with a tone for 3 min. Then contextual fear extinction was tested by placing the mice for 5 min in the same context used for CS-US conditioning for 5 consecutive days (Days 2-6) with no tone or shock presentation. Freezing was recorded during each test and analyzed using Image FZC 2.22 sr2 software (O’Hara & Co Ltd., Japan).

### Forced swim test

The forced swim test was performed as described by Inoue et al. 2008^94^. Briefly, mice were placed in water-filled cylinder for 10 min. Immobility was recorded and analyzed starting from the third minute using Time software (O’Hara & Co Ltd., Japan).

### Elevated-plus maze test

The maze consisted of two open and two closed arms with dimensions (25 cm length * 5 cm width), which were elevated 50 cm above the floor. Mice were placed in the center region facing one of the open arms and allowed to move freely for 10 min. Time spent in open arms was recorded and analyzed using Time EPC software (O’Hara & Co Ltd., Japan).

### Open field test

Mice were placed in the center of an open field arena with dimensions (50 * 50 * 33.3 cm; width, depth and height) and 100 lux illumination intensity and allowed to freely move for 15 min. Movement traces, speed, distance travelled, and time spent in the center of the open field were recorded and analyzed using Time OFCR software (O’Hara & Co Ltd., Japan).

### Tail suspension test

Mice were suspended by their tails for 6 min. Immobility duration was recorded and analyzed using Time software (O’Hara & Co Ltd., Japan).

### Adeno-Associated Virus (AAV) production

AAV serotype 9 expressing mouse *Tob* under control of the human synapsin promoter (hSyn) was generated as described by Kudo et al., 2020^95^. Briefly, pAAV2-hSyn-mTob was generated from pAAV2-hSyn-EGFP (Addgene, USA) by replacing the EGFP sequence with mouse *Tob* coding sequence. AAV-293 cells (Agilent, USA) were transfected with AAV-rep2/cap9 expression plasmids, adenovirus helper plasmids, and AAV-vector plasmids to generate AAV9-hSyn-mTob.

### Stereotactic surgery for viral injection

*Tob*^fl/fl^ mice were bilaterally injected with Cre-expressing adeno-associated virus AAV1.hSyn.Cre.WPRE.hGH (105553-AAV1, Addgene) to generate hippocampus-specific KO mice. *Tob*-WT and KO mice were injected with AAV to express mouse TOB AAV9.hSyn.mTob.WPRE.hGH for rescue experiments. Stereotaxic surgical procedures were performed as described by Augustinaite and Kuhn, 2020^96^. Briefly, mice were anesthetized using an intraperitoneal injection of a mixture of Medetomidine (0.3 microgram/g), Midazolam (5 μg/g) and butorphanol (5 μg/g). Additionally, a non-steroidal anti-inflammatory, Carpofen (7.5 μg/g), was injected by the end of the surgery. Mice were fixed on a stereotaxic frame and head hair was shaved. A 2% lidocaine solution was applied to the shaved skin and left for 2 min. Iodine was applied gently over the skin as an antiseptic. A midline incision was made, and skin was retracted, and the skull was exposed. After drying the surface, the bregma was detected. A micromanipulator was used to slowly move the injection needle to the target injection site. A dental drill was used to drill a small hole, until the surface of the brain appeared. A needle with viral solutions of around 300 nL was slowly advanced into the hole until it touched the brain surface, and slowly lowered to the target coordinates. Injection was done over 2 min and thereafter, the needle was left in place for 5 min before slowly retracting it. Coordinates used for the CA1 region of the hippocampus were tested and optimized as 1.6 mm posterior, 1.5 mm medio-lateral and 1.6 mm ventral to the bregma.

### Immunohistochemistry

Immunohistochemical staining was performed as described by Matsuura et al., 2021^97^. Antibodies used were Anti-Cre RpAb (Cell Signaling, USA) and Alexa Flour 488 Goat Anti-rabbit IgG (Invitrogen, USA).

### Statistical analysis

All data are presented as means ± SEMs. T-tests, Mann-Whitney U test, one-way ANOVA, and two-way ANOVA were used as described in figure legends. Multiple testing following ANOVA was corrected using Bonferroni or Dunnet’s post-hoc tests. GraphPad prism 9 was used to perform all statistical analyses.

## Acknowledgments

This work was financially supported by Okinawa Institute of Science and Technology Graduate University and JSPS Kakenhi Grant number 18J20551. We thank Prof. Yoko Yazaki-Sugiyama and Dr. Yuichi Morohashi for their support in generating Tob AAV viral vectors. pAAV-hSyn-EGFP was a gift from Bryan Roth (Addgene plasmid # 50465), pAdDeltaF6, pAAV2/9n, pENN.AAV.hSyn.Cre.WPRE.hGH were gifts from James M. Wilson (Addgene plasmid # 112867, 112865, Addgene viral prep # 105553-AAV1). Illustrative figures were created with Biorender.com. We are grateful for the help and support provided by the animal resources, high-performance computing, and sequencing sections of the Research Support Division at Okinawa Institute of Science and Technology Graduate University.

## Author Contributions

MY and TY conceived the idea and coordinated the study. MY performed the behavioral experiments, molecular experiments and bioinformatic analyses. HH performed fMRI. EL performed electrophysiological recording. ME and BK performed stereotaxic surgery and viral injections. YK performed elevated-plus maze and provided support for behavioral analysis. HK and KN generated Tob^fl/+^ mice. MY, EL, HH, ME and TY participated in manuscript writing. All authors revised and approved the final version of manuscript.

## Supplementary information

Supplementary information is available at journal’s website.

