## Supplementary information for "TOB is an effector of the hippocampus-mediated acute stress response"

### Supplementary materials and methods:

#### Generation of *Tob* conditional knockout mice

Tob$^{flox/+}$mice (Accession No. CDB0044E; http://www2.clst.riken.jp/arg/mutant%20mice%20list.html ) were generated by single-strand oligodeoxynucleotide (ssODN) knock-in with CRISPR/Cas9-mediated genome editing using C57BL/6N zygotes, in which the exon2 of *Tob* gene was flanked by loxP sites. The two loxP sites were introduced one-by-one as follows. In the first step electroporation, the 3'-loxP site was introduced into wild type zygotes and the resulting F0 mice were genotyped to identify the 3'-loxP site. For the second step electroporation, the 5'-loxP site was introduced into the zygotes carrying the 3'-loxP site which were generated by in vitro fertilization between wild type females and the 3'-loxP males. For electroporation, the zygotes were transferred into Opti-MEM I (ThermoFisher Scientific) containing a crRNA (25ng/ul), tracrRNA (100ng/ul), an ssODNs consisting of a loxP site and a restriction enzyme recognition site (EcoRI) (400ng/ul) and Cas9 protein (250ng/ul) (ThermoFihser Scientific). CUY21 EDIT II and LF501PT1-10 platinum plate electro code (BEX Co. Ltd.) were used and electroporation conditions were 30 V (3 msec ON + 97msec OFF) x 7 times as described by Hashimoto *et al.* (2015)[1]. After electroporation, the zygotes were transferred into pseudopregnant ICR female mice. gRNA sites were designed by using the Zhang lab website (Nat Biotechnol. 2013 Sep;31(9):827-32.) and the crRNAs, tracrRNA were as follows; 5'-crRNA (5'-AGA CCG GUG UUU GCC AGA GCg uuu uag agc uau gcu guu uug-3'), 3'-crRNA (5'- AGA CCG GTG UUU GCC AGA GCg uuu uag agc uau gcu guu uug-3'), tracrRNA (5'-AAA CAG CAU AGC AAG UUA AAA UAA GGC UAG UCC GUU AUC AAC UUG AAA AAG UGG CAC CGA GUC GGU GCU-3'), 5'ssODN (5'-GGA AAC TAA TGA CCA ACT TGT AAA AGT GGA GTG TAC GGG TGA ATT CAT AAC TTC GTA TAA TGT ATG CTA TAC GAA GTT ATA GGT GGG TGG ATG AGT GTG CAG TAC AGT TAG TAA ACA CTG-3') and 3'ssODN (5'-GAA GGC AGG GCT CTG CTG TTT GTC AGA CCG GTG TTT GCC AGA ATT CAT AAC TTC GTA TAA TGT ATG CTA TAC GAA GTT ATG AGC TGG GGT GGT GCA TAG TAG AGT GTT TGC CTC AAA AGC-3') (Facmac Co., Ltd). The loxP sites were confirmed by PCR and sequencing. The PCR was performed following primers; 5'FW (5'-TGA GAG CCC TTG GCA TGG-3') and 5'REV (5'-ATA CCA CTT CCC AGC AGG-3') (WT: 361 bp, 5'-loxP: 401 bp) for 5'-loxP site, and 3'FW (5'-GGA ATA ATG GAA GGC AGG-3') and 3'REV (5'-CCT CCT ATC ACC TGG CTC-3') for 3'-loxP site (WT: 190 bp, 3'-loxP: 230 bp), respectively (Fig. S4A). The PCR product sizes after EcoRI digestion were 5'-loxP (258 bp and 143 bp) and 3'-loxP (50 bp and 180 bp). The protocol for animal experimentation was approved by the Institutional Animal Care and Use Committee (IACUC) of RIKEN Kobe Branch, and all the animal experiments were conducted in accordance with the institutional guideline.

#### Resting state functional magnetic resonance imaging (rs-fMRI)

*Tob*-KO mice (12 weeks; mean body weight = 25.86, std = 1.49, n=8) and *Tob*-WT (12 weeks; body weight = 26.9, std = 0.70, n=8) mice were introduced to perform MRI experiments. The difference in the body weight between the two groups showed no statistical significance (p-value = 0.16).

In order to minimize head movements inside MRI, mice undergone surgery to mount head-fixation bar on the skull of each mouse after the age of 8 weeks as described in details by Takata et al., 2018[2], Abe et al., 2020[3] and Yoshida et al., 2016[4]. Briefly, an anesthetic mixture (0.3 mg/kg medetomidine, 4 mg/kg midazolam, and 5 mg/kg butorphanol) was intraperitoneally injected into mice. Mice were subsequently fixed to a stereotaxic apparatus (SM-15, Narishige Scientific Instrument, Japan). The skull surface was exposed to mount a custom-made acrylic head-fixation bar (3 x 3 x 27 mm3) using dental cement (Super-Bond

C&B, Sun Medical, Shiga, Japan). After their body weights were recovered, mice were acclimated to a mock MRI environment for 2 hours/day for 7 days. During the acclimation, MRI scanner sounds recorded in actual fMRI experiments were delivered through speakers.
MRI images were acquired with an 11.7 T MRI scanner for small animals (Biospec 117/11 system, Bruker Biospin, Germany) with a cryogenic quadrature RF surface probe (Cryoprobe, Bruker BioSpin AG, Switzerland). Respiration rate and rectal temperature (37°C) were monitored during scanning and body temperature was maintained at 37°C using an MR-compatible, feedback-controlled water heating system.
Structural (anatomical) images for spatial normalization were obtained using a 3D-multislice rapid acquisition with relaxation enhancement (RARE) sequence with the following parameters: TR/TE = 5449/15.7 ms, FOV=13.5 × 13.5 × 12.5 mm3, matrix = 94 × 94, resolution = 0.14 × 0.14 × 0.3 mm3, 41 coronal slices, scan time = 2 min.
Gradient echo (GE)-EPI sequence was performed in the same brain locations for resting state fMRI with the following parameters: repetition time (TR)/echo time (TE) = 2000/14 ms, segments=1, FOV = 13.5 × 13.5 mm2, matrix = 90 × 90, resolution = 0.15 x 0.15 x 0.3 mm3, 41 coronal slices, repetition number = 300, total imaging time = 10 min.

**Preprocessing and de-nosing of fMRI data**

First five functional images in each run were removed from analyses in order to prevent contamination of initial imaging artifacts due to motion and non-steady-state image quality. 10-fold magnification of images was done to process images with Statistical Parametric Mapping 12 (SPM12; Wellcome Trust Centre for Neuroimaging, UK), which is designed to process human-size brain images. Therefore, voxel size was 1.5x1.5x3mm in the analysis steps. Pre-processing including motion correction, realignment, co-registration, normalization to a C57BL6/J template, and spatial smoothing (kernel with 3x3x6mm) were executed by Statistical Parametric Mapping (SPM12). Since each slice of 3D brain image was taken at different timing, slice timing correction was applied to correct slice-timing difference by temporal interpolation. Next, the realignment of 3D brain images was conducted to correct motion-related changes of brain position. Then, the co-registration step was executed to overlay functional T2* images to structural T2 images and save coordination change of T2* images for the normalization step. In the normalization step, T2 structural image was first warped to fit the average C57BL6/J template[5], and T2* functional images were further warped to the template using the co-registered coordination change of the images. In the spatial smoothing, voxel signals were spatially smoothed with a gaussian kernel (3x3x6mm).
In the de-noising process, linear de-trend filtering, temporal filtering (0.01-0.08 Hz), 6 motion regressions, signal regression of grey matter (GM), white matter (WM), and cerebrospinal fluid (CSF), de-spiking, and motion scrubbing were executed to reduce false-positive functional connectivity[6] with functional connectivity toolbox (CONN17, available at: https://www.nitrc.org/projects/conn)). Linear de-trend filtering was applied to remove linear accumulation of imaging noise. Temporal filtering was done with 0.01-0.08(Hz) frequency range. Six Motion-related artifacts in T2* signal such as x, y, z coordinates and pitch, raw, yaw rotations were regressed out in the 6 movement regressions. Average signals of WM and CSF were regressed out to reduce the potential influence of non-physiological signals, and average GM was further regressed out to conduct global regression. Motion-related artifacts were further regressed out with a squashing function in the de-spiking step. Prior to the analyses, motion scrubbing was finally executed to remove signal outliers (z-score > 5).

**Electrophysiological recording**

Mice, with age of P18-P30, were deeply anesthetized with $CO_2$ and euthanized by decapitation. The brain was quickly removed and placed in the chilled (0–4°C) artificial cerebrospinal fluid

(ACSF) containing (in mM) 126 NaCl, 2.5 KCl, 1 $CaCl_2$, 1 $MgSO_4$, 1.25 $NaH_2PO_4$, 26 $NaHCO_3$, and 20 glucose, bubbled with 95% $O_2$ and 5% $CO_2$ (pH 7.4). 250 μm thick parasagittal hippocampal slices were prepared using a vibratome slicer (VT-1200S, Lecia, Germany). Hippocampal slices were allowed to recover first at 32ºC for 30 min and then at room temperature for another 30 min before recordings. Whole-cell recordings were made from visually identified somata of hippocampal pyramidal neurons at 32°C using an upright microscope (BX50WI, Olympus, Japan). Two intracellular solutions were used with the following compositions (in mM): (1) 132.5 Cs-gluconate, 17.5 CsCl, 2 $MaCl_2$, 10 HEPES, 10 BAPTA-$K_4$, 4 ATP, 0.4 GTP (pH 7.3, adjusted with CsOH) for recording excitatory postsynaptic currents (EPSCs), (2) 124 CsCl, 1 $CaCl_2$, 4.6 $MgCl_2$, 10 HEPES, 10 BAPTA-$K_4$, 4 ATP, 0.4 GTP (pH 7.3, adjusted with CsOH) for recording inhibitory postsynaptic currents (IPSCs). Picrotoxin (100 μM) was added to block GABA receptor-mediated inhibitory currents for recording EPSCs, and NBQX (10 μM) and D-AP5 (50 μM) were included to block AMPA receptor-mediated and NMDA receptor-mediated excitatory synaptic currents, respectively for recording IPSCs. AMPAR-mediated EPSCs recordings were conducted with D-AP5 (50 μM) in the ACSF to isolate NMDAR responses and 0.1 mM Spermine added to the internal solution. NMDAR-mediated ESPCs recordings were conducted in the presence of NBQX (10 μM) to isolate the AMPAR responses. Recordings of synaptic activity were made using a bipolar electrode to deliver stimuli (0.05 Hz) to the Schaffer collateral pathway to evoke postsynaptic currents in CA1 pyramidal neurons.
mEPSCs and mIPSCs recordings were performed with CA1 pyramidal neurons voltage clamped at -70mV. 1 μM tetrodotoxin (TTX, Tocris) was added in ACSF to block action potential evoked responses; 100-200 events from each sweep were used for data analyses.
To assess input-output relationship, stimuli of increasing intensities (1-10 μA) were applied every 10s. AMPAR-mediated currents were recorded at a membrane holding potential at -70 mV, while NMDAR-mediated currents were recorded at a holding potential of +40 mV. Ten traces were collected and averaged at each stimulus. Summary graphs were constructed by normalizing all values to the eEPSC at 1 μA stimulus intensity. To record current-voltage relationship, peak amplitude of eEPSCs were measured at holding potential from -80mV to +60mV. Ten traces were collected and averaged at each holding potential. Summary graphs were constructed by normalizing all values to the eEPSC at -80 mV. Rectification index was calculated by current ratio of EPSCs between +40 and -40 mV.

AMPAR-mediated Paired-pulse ratio (PPR) experiments were performed at −70 mV holding potential. The amplitude of the second EPSC was measured relative to the amplitude of the first EPSC. PPR was measured at six inter-stimulus intervals, 10 ms, 30 ms, 50 ms, 100 ms, 200 ms and 300 ms. Twenty traces were collected for each inter-stimulus interval.

To measure the excitatory/inhibitory (E / I) ratio, AMPA receptor-mediated currents evoked were first recorded at a holding potential of -60 mV. Neurons were subsequently depolarized to 0 mV, and $GABA_A$ receptor-mediated synaptic currents were recorded. The inhibition/excitation ratio was calculated by dividing the peak of AMPPA receptor-mediated current by the peak of $GABA_A$ receptor-mediated current. All recordings were digitized at 10 kHz and filtered at 3 kHz. Recordings were monitored using a Multiclamp 700B amplifier (Molecular Devices), and a Digidata 1440A digitizer (Molecular Devices). Online data acquisition and offline data analysis was performed using Clampex and Clampfit version 10.7 (Molecular Device) or Mini analysis Program (ver. 6.0.3, Synaptosoft Inc.). Statistical significance of data was evaluated using a Unpaired two-tailed Student's *t*-test and Mann-Whitney U test. For cumulative probability plots, a Kolmogorov–Smirnov test with a significance of $p < 0.05$ was used. For comparing the slopes of linear curves, simple linear regression was used. All recordings and analysis were done with experimenter blind to genotype.

**Supplementary Figures:**

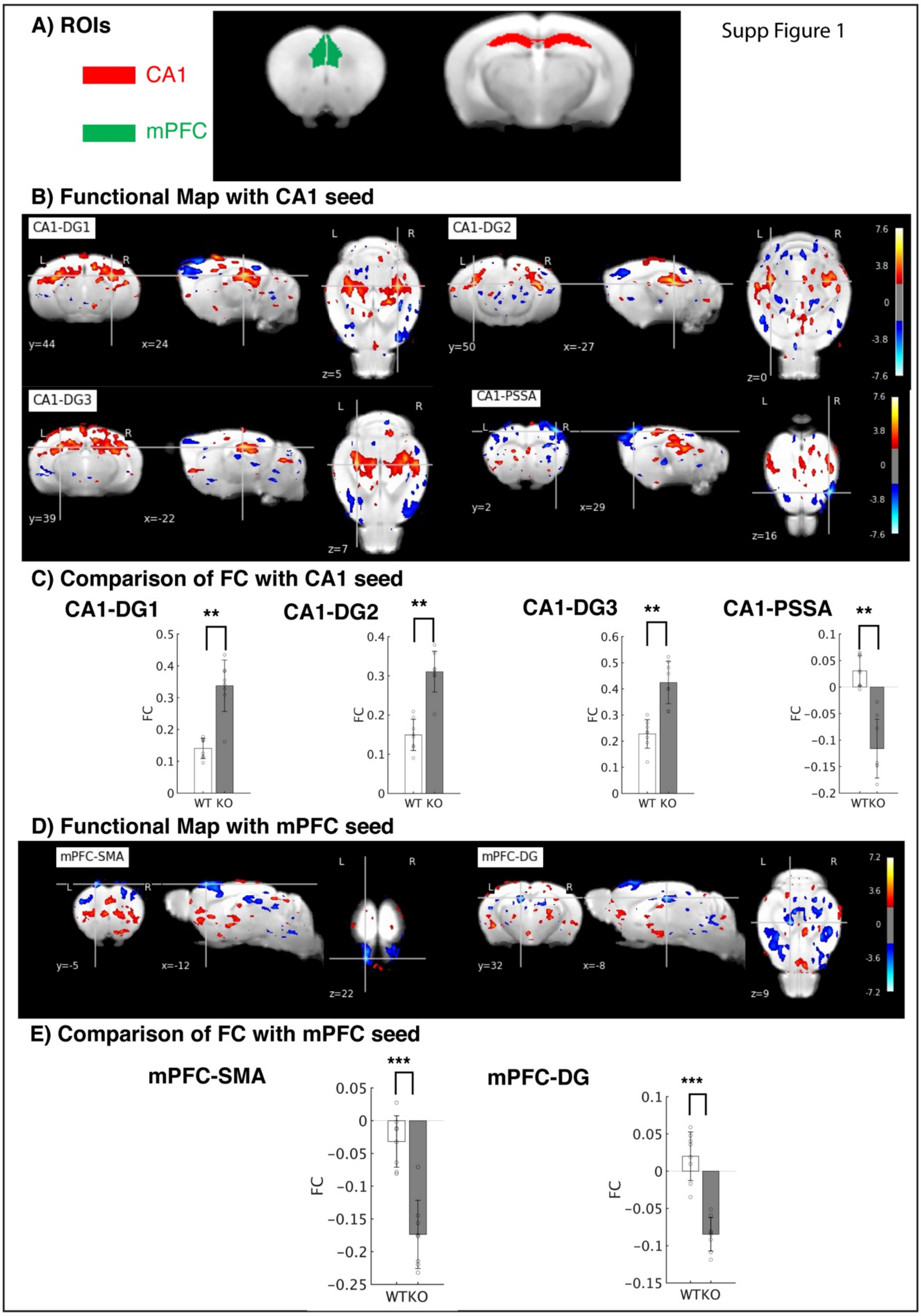


**Fig. S1: Effect of *Tob* deletion on functional connectivity between different brain regions**. A) CA1 and mPFC seeds. The seed regions were extracted from the Allen Brain Atlas (ABA). B) Functional map with the CA1 seed between *Tob*-KO (n=8) and WT mice (n=8). Functional map of the CA1 seed with subfields of the DG, DG1-3, and PSSA was visualized (thresholded with |T| > 2, uncorrected). C) Comparison of FC of the CA1 seed. FCs of the CA1 seed were calculated with cluster-corrected regions including DG1-3 and PSSA. The FCs between *Tob*-KO and WT mice in CA1-DG revealed statistically positive significance while FC in CA1-PSSA revealed statistically negative significance. ** $p<0.01$. D) Functional map with the mPFC seed between *Tob*-KO (n=8) and WT mice (n=8). Functional map of the mPFC seed with the SMA and right DG was visualized (thresholded with |T| > 2, uncorrected). E) Comparison of FC of the mPFC seed. FCs of the mPFC seed were calculated with cluster-corrected regions including SMA and right DG. The FCs between *Tob*-KO and WT mice in mPFC-SMA and -DG revealed statistically negative significance. *** $p<0.001$. mPFC: medial prefrontal cortex; CA1: cornu ammonis 1; CA2: cornu ammonis 2; CA3: cornu ammonis 3; DG: dentate gyrus; PSSA: primary somatosensory Area.

### NMDAR-mediated neurotransmission

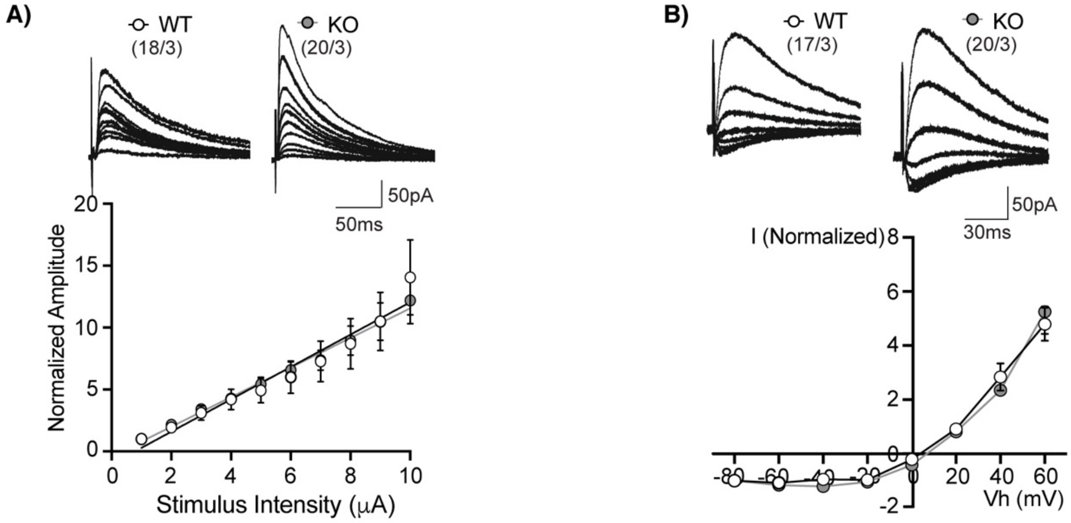


**Fig. S2**: ***Tob* deletion has no effect on NMDA-mediated synaptic transmission at CA3-CA1 synapses in the hippocampus.** A) Sample traces (upper panel) and summary plots for input-output relationship of NMDA receptor-mediated responses recorded from WT (open circles) and *Tob*-KO (grey circles) mice. Scale bars, 50 pA and 50 ms. B) Sample traces (upper panel) and summary plots for I-V curve of NMDA receptor-mediated responses recorded from WT (open circles) and *Tob*-KO (grey circles) mice. Scale bars, 50 pA and 30 ms.

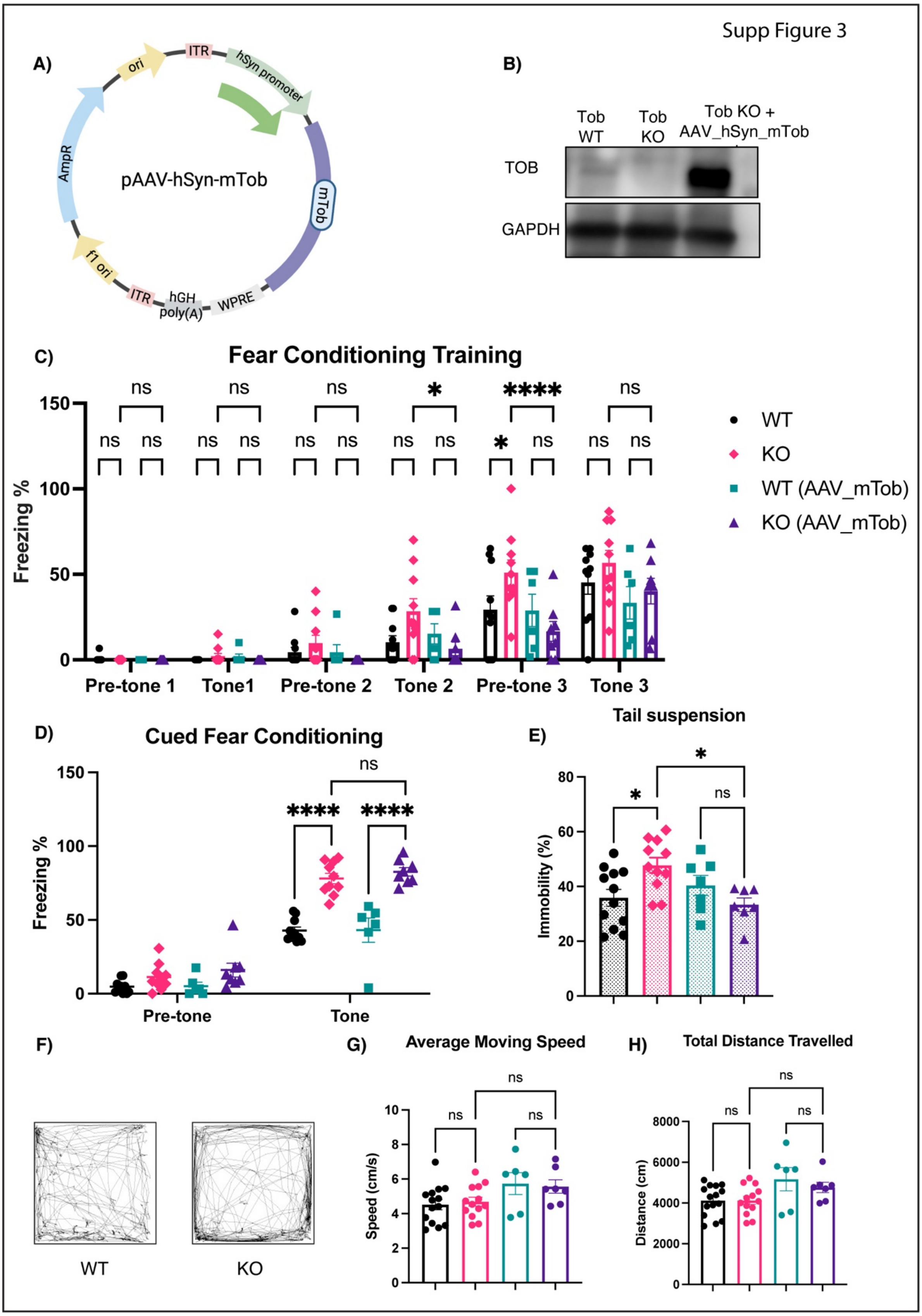


**Fig. S3: Behavior in *Tob*-KO mice and effect of overexpressing TOB using AAV_hSyn_mTob in the hippocampus**. A) Schematic diagram for the structure of plasmid used to generate AAV_hSyn_mTob. B) Immunoblot showing TOB expression after

AAV_mTob injection in *Tob*-KO mice hippocampi. (C-D) Fear Conditioning. C) Freezing during fear conditioning training measure at three pre-tone and tone (with electric shock) presentation time periods. *Tob*-KO mice showed increased freezing at pre-tone 3 when compared to WT mice (KO vs WT at pre-tone 3 $p=0.0106$). There were no significant differences in freezing between *Tob*-WT, KO and KO(AAV_mTob) throughout the training session. D) Cued fear conditioning represented as freezing when tone is presented. There was no significant difference between *Tob*-KO and KO(AAV_mTob) mice, but they showed increased cued fear freezing when compared to wild-type controls ($F_{3,32}=22.28$ for genotype effect $p<0.0001$; WT vs KO at tone $p<0.0001$; WT(AAV_mTob) vs KO(AAV_mTob) at tone $p<0.0001$; KO vs KO(AAV_mTob) $p>0.9999$). Two-way ANOVA followed by Bonferoni's *post-hoc* test for multiple comparisons test. E) Tail suspension test represented as immobility. Immobility was higher in *Tob*-KO mice, which was rescued by AAV-mediated TOB expression ($F_{3,33}=4.123$ $p=0.0137$; WT vs KO $p=0.0332$; KO vs KO(AAV_mTob) $p=0.0295$; WT(AAV_mTob) vs KO(AAV_mTob) $p>0.9999$). (F-H) Open field test. F) Representative figure for *Tob*-WT and KO mice movement traces in open field test. G) *Tob*-KO mice show normal moving speed during open field test. H) *Tob*-KO mice show normal movement presented as distance travelled during open field test. One-way ANOVA followed by Bonferoni's *post-hoc* test for multiple comparisons test. All values represent means±SEM. ns non-significant, * $p<0.05$, ****$p<0.0001$

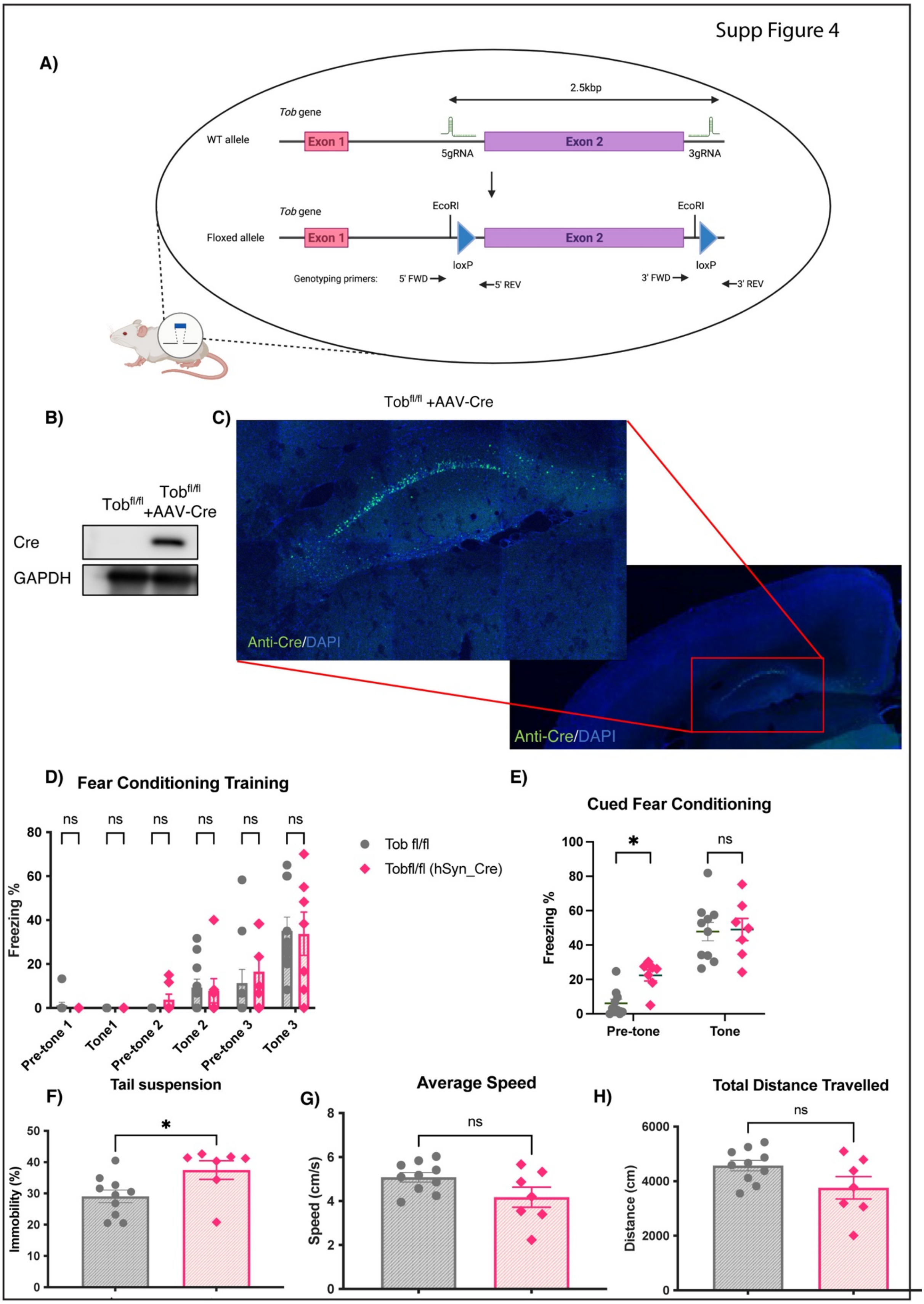


**Fig. S4: Behavior of hippocampus-specific knockout mice (hsTob KO) mice.** A) Schematic diagram showing the procedure for generating floxed *Tob* mice (Tob$^{fl/fl}$) through insertion of two LoxP sequences flanking the coding exon (exon2) of *Tob* gene. B) Immunoblot showing

Cre expression after AAV_Cre injection in $Tob^{fl/fl}$ mice hippocampi. C) Confocal brain images of $Tob^{fl/fl}$ after AAV_Cre injection. Images on the right depict outlined areas to shows hippocappus-specific Cre expression (Green). Nuclei are labeled with DAPI (blue). Fear Conditioning (D-E). D) Hippocampal *Tob* deletion had no effect on freezing levels during fear conditioning ($F_{1,15}$=0.06450 for genotype effect p=0.8030). E) Cued fear conditioning represented as freezing when tone is presented. hsTob KO mice showed similar freezing levels after tone presentation to $Tob^{fl/fl}$ mice ($F_{1,18}$=3.031 for genotype effect at p=0.0988; $Tob^{fl/fl}$ vs $Tob^{fl/fl}$(hSyn_Cre) at tone p>0.9999, at pre-tone p=0.0297). Two-way ANOVA followed by Bonferoni's *post-hoc* test for multiple comparisons test. F) Tail suspension test represented as immobility. Immobility was higher in hsTob KO mice (p=0.0282). (G-H) Open field test. G) hsTob KO mice show normal moving speed during open field test. H) hsTob KO mice show normal movement presented as distance travelled during open field test. Unpaired t-test. All values represent means±SEM. ns non-significant, * p<0.05.

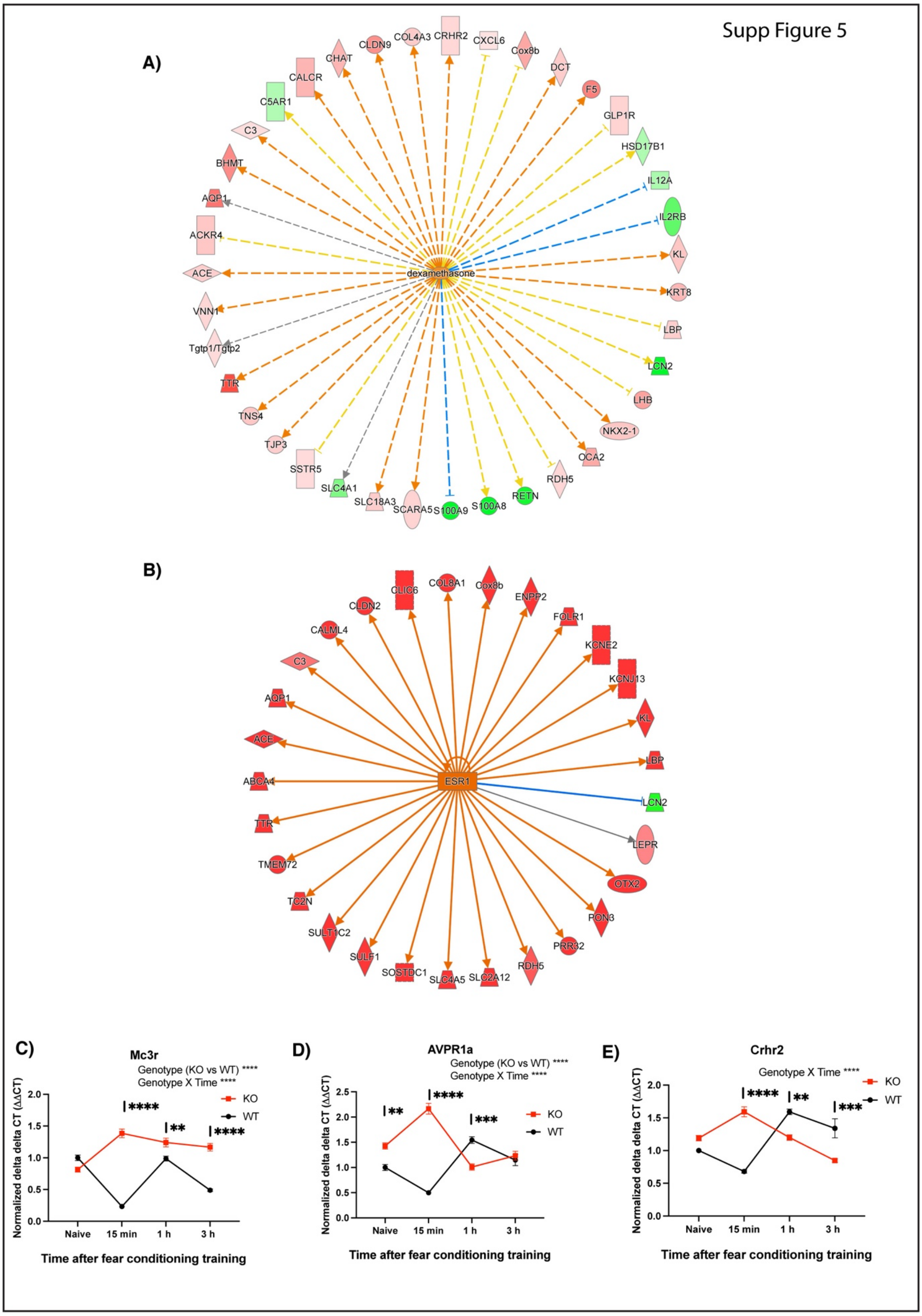


**Fig. S5: Pathway analysis for differentially expressed genes in hippocampus of *Tob*-KO mice at 15 minutes post fear conditioning**. A) Differential expression for 39 genes (30 upregulated and 9 downregulated) suggests activation of dexamethasone upstream pathway.

B) Differential expression for 29 genes (28 upregulated and 1 downregulated). C) Quantitative real-time PCR for *Mc3r*, *Avpr1a* and *Crhr2* mRNAs which showed increased expression at 15 min post-fear conditioning (Mc3r, $F_{1,16}$=191.0 for genotype effect at p<0.0001, WT vs KO at 15 min. p<0.0001; Avpr1a, $F_{1,16}$=55.38 for genotype effect at p<0.0001, WT vs KO at 15 min. p<0.0001; Crhr2 $F_{1,16}$=1.317 for genotype effect at p=0.2681, WT vs KO at 15 min. p<0.0001). Two-way ANOVA followed by Bonferoni's *post-hoc* test for multiple comparisons test. All values represent means±SEM. ** p<0.01, *** p<0.001, ****p<0.0001
